# To cure or not to cure: consequences of immunological interactions in CML treatment

**DOI:** 10.1101/494575

**Authors:** Artur César Fassoni, Ingo Roeder, Ingmar Glauche

**Affiliations:** Artur César Fassoni: Instituto de Matemática e Computação, Universidade Federal de Itajubá, Itajubá, Brazil; Ingo Roeder: Institute for Medical Informatics and Biometry, Carl Gustav Carus Faculty of Medicine, Technische Universität Dresden, Dresden, Germany. National Center for Tumor Diseases (NCT), Partner Site Dresden, Germany; Ingmar Glauche: Institute for Medical Informatics and Biometry, Carl Gustav Carus Faculty of Medicine, Technische Universität Dresden, Dresden, Germany

**Keywords:** Mathematical Modeling, Ordinary Differential Equations, Bifurcations, Chronic Myeloid Leukemia, Immunological Control, Treatment-Free Remission

## Abstract

Recent clinical findings in Chronic Myeloid Leukemia (CML) patients suggest that the number and function of immune effector cells are modulated by Tyrosine Kinase Inhibitors (TKI) treatment. There is further evidence that the success or failure of treatment cessation at least partly depends on the patient’s immunological constitution. Here, we propose a general ODE model to functionally describe the interactions between immune effector cells with leukemic cells during the TKI treatment of CML. In total, we consider 20 different sub-models, which assume different functional interactions between immune effector and leukemic cells. We show that quantitative criteria, which are purely based on the quality of model fitting, are not able to identify optimal models. On the other hand, the application of qualitative criteria based on a dynamical system framework allowed us to identify nine of those models as more suitable than the others to describe clinically observed patterns and, thereby, to derive conclusion about the underlying mechanisms. Additionally, including aspects of early CML onset, we can demonstrate that certain critical parameters, such as the strength of immune response or leukemia proliferation rate, need to change during CML growth prior to diagnosis, leading to bifurcations that alter the attractor landscape. Finally, we show that the crucial parameters determining the outcome of treatment cessation are not identifiable with tumor load data only, thereby highlighting the need to measure immune cell number and function to properly derive mathematical models with predictive power.

*MSC Classification*: 92B05, 37N25, 34C60, 37G35

## 1 Introduction

Chronic Myeloid Leukemia (CML) is a hematological cancer characterized by the clonal expansion of myeloid cells in the bone marrow [Hoffbrand et al., 2001]. In the majority of cases, CML results from a single genetic alteration in hematopoietic stem cells, namely a translocation that involves the BCR gene in chromosome 22 and ABL1 gene in chromosome 9 [Chereda and Melo, 2015]. The resulting BCR-ABL1 fusion gene is continuously activated, thereby triggering a cascade of proteins that deregulate the cell cycle and accelerate cell division. Due to their unregulated growth and their limited differentiation, leukemic cells accumulate in bone marrow and outcompete the normal blood cells [Hehlmann et al., 2007].

In the last decades, the treatment of CML was revolutionized by the advent of targeted therapy with Tyrosine Kinase Inhibitors (TKIs), a family of drugs which targets the BCR-ABL1 fusion protein and thereby preferentially affects mutated cells [Druker et al., 2006, Hochhaus et al., 2017]. Normal cells remain largely unaffected leading to less side effects and better disease control than standard cytotoxic chemotherapy. Thus, CML treatment with TKIs was one of the first successful examples of targeted therapy, and a formerly lethal cancer was turned into a controllable disease in which patients have close-to-normal life expectancy [Jabbour and Kantarjian, 2016, Bower et al., 2016]. Although the therapy with TKIs achieved high long-term survival rates, only few patients with very good treatment response qualify for treatment cessation studies, which reproducibly report treatment free remission (TFR) in only 50 - 60 % of cases [Mahon et al., 2010, Rousselot et al., 2014, Rea et al., 2017b, Saussele et al., 2018]. For all other patients, permanent disease control requires continuing and a potentially life-long therapy. Ongoing efforts seek to understand the processes accounting for residual disease control and for defining criteria for safe treatment discontinuation [Saussele et al., 2016].

Multiple studies pointed towards a critical role of the immune system as an important player in the control of residual disease after TKI stop [Ilander et al., 2017, Hughes et al., 2017, Rea et al., 2017a, Sopper et al., 2016, Schütz et al., 2017, Kumagai et al., 2018]. It has been speculated that, once the treatment has removed CML load below a certain threshold, the immune cells are further able to limit the growth of the residual tumor, leading to sustainable treatment-free remission, or, in some cases, even to the complete eradication of leukemic cells [Hughes and Yong, 2017]. However, there are few clinical measurable indicators for quantifying such immune control and the mechanisms underlying these interactions are still controversial [Hughes and Yong, 2017]. Specially the lack of quantitative information on interactions of the immune system with CML cells makes it difficult to derive robust criteria for conceptual model approaches and for the prediction of TKI cessation scenarios.

Mathematical models provide an elegant means to study different modes of CML-immune interactions and their functional consequences. Especially the well-defined phenotype and the accessibility of kinetic data on treatment responses made CML a showcase example for the application of mathematical modelling approaches [Michor et al., 2005, Roeder et al., 2006, Dingli et al., 2010, Komarova and Wodarz, 2009, Stein et al., 2011]. Several models provided better understanding of the treatment dynamics by dissecting the mechanisms underlying the observed treatment response in CML patients. Some considered the role of the immune system on tumor progression [Kim et al., 2008, Wodarz, 2010, Clapp et al., 2015, Besse et al., 2018], but different hypothesis on the interactions between the leukemic and immune cells were assumed. For instance, a functional form describing an immune recruitment with a saturation for large numbers of tumor cells was mechanistically derived in [Kuznetsov et al., 1994], and a functional form describing an optimal immune window for both the immune response and recruitment were proposed in [Clapp et al., 2015]. For comprehensive reviews on models for interaction between tumor and immune response, see [Eftimie et al., 2011] and [Clapp and Levy, 2015].

Using and integrating the knowledge gained from these previous works, we consider the most frequent and plausible assumptions occurring in those different models, and compile them in a systematic dynamical analysis of a generalized CML-immune model. Based on the current clinical presentation of CML under TKI treatment, we define several criteria that a suitable model should satisfy. Technically, we consider a general mathematical model based on ordinary differential equations (ODEs), to qualitatively describe the immune control of residual leukemic cells prior, during and after TKI therapy. The general model encompasses a set 20 different sub-models, each of which considers different, possible assumptions on the two main CML-immune interactions: (i) the targeting of CML cells by immune cells and (ii) the stimulation or inhibition of immune cells due to the presence of CML cells (referred to as recruitment). We analyze each sub-model in detail and qualitatively compare the features of each sub-model with available data from TKI cessation studies [Mahon et al., 2010, Rousselot et al., 2014].

Our approach allows comparing the consequences of combining different model assumptions about the immunological control of CML cells by the immune system as well as opposing effects of the leukemia on the immune system. We investigate, from a dynamical systems point of view, the possible routes from CML onset until diagnosis and then treatment and treatment-cessation outcomes. This model analysis provides insights into the mechanisms involved in the immunological control of CML after TKI cessation.

## 2 Mathematical Model

We apply an ODE model to describe the dynamics of CML progression and its interactions with the immune system. This model extends a recently published model in the context of TKI dose reduction [Fassoni et al., 2018] by including a compartment of immune effector cells. The model considers quiescent and proliferating leukemic stem cells (LSCs) as well as immune cells, denoted by *X*, *Y* and *Z*, respectively. For reasons of simplicity, the immune cell population is considered as a population that comprises all immune effector cells responsible for the immune response; such as NK cells and CD8+ T cells. A scheme of the model is depicted in Figure 1. The model is formally written as the following ODEs system:

**Figure 1:**
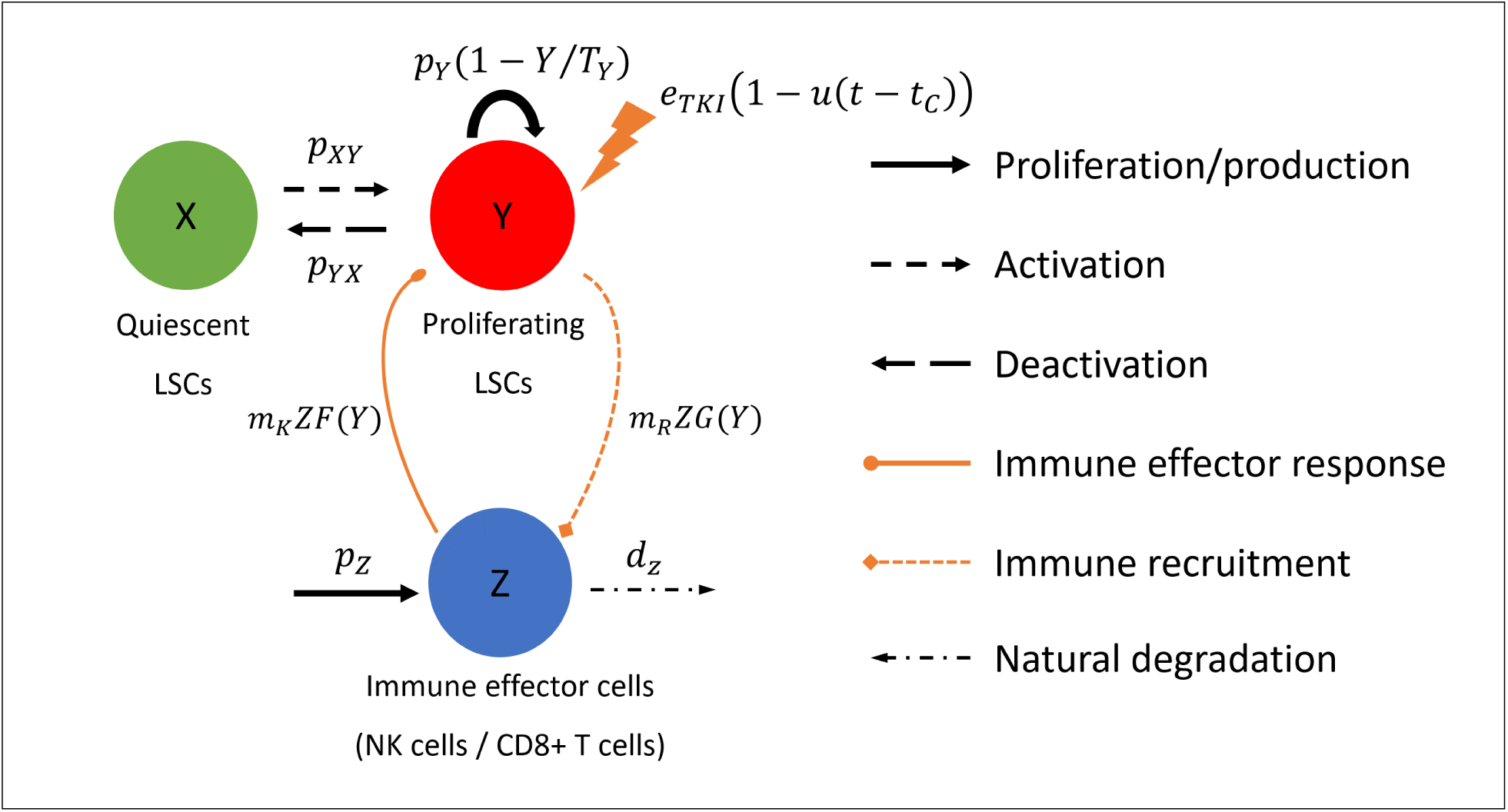
Schematic model representation with three cell types: quiescent (*X*, green) and proliferating (*Y*, red, turnover with rate *p*_*Y*_ (1 − *Y/T*_*Y*_)) leukemic stem cells, and immune effector cells (*Z*, blue, generated with rate *p*_*Z*_, decaying with rate *d*_*Z*_). The model assumes (i) mechanisms of activation/deactivation of quiescent/proliferating LSCs with rates *p*_*XY*_ and *p*_*YX*_ ; (ii) a cytotoxic effect of TKI on proliferating LSCs with intensity *e*_*TKI*_, applied from time *t* = 0 until the cessation time *t* = *t*_*C*_; (iii) an immune response (targeting) on proliferating LSCs with intensity *m*_*K*_*ZF*(*Y*) (per capita rate *m*_*K*_, modulated by *F*(*Y*)); and (iv) an immune recruitment due to the encounter of immune effector with proliferating LSCs, with intensity *m*_*R*_*ZG*(*Y*) (per capita rate *m*_*R*_, modulated by *G*(*Y*)).

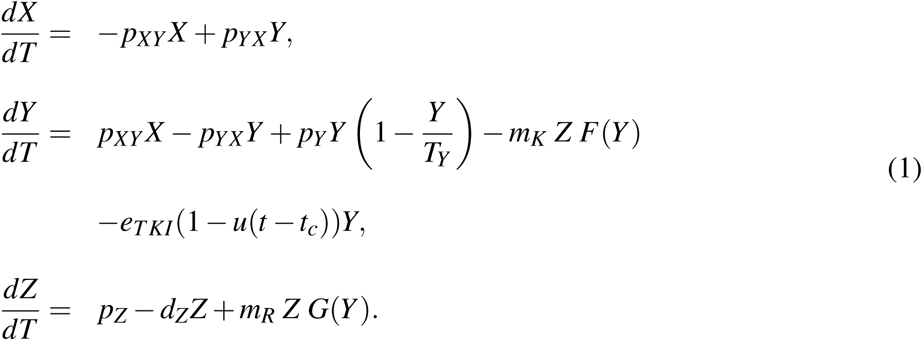

The model assumes that quiescent LSCs do not proliferate and enter the proliferating compartment with activation rate *p*_*XY*_. This compartment of quiescent LSCs is assumed to be insensitive to TKI [Holyoake and Vetrie, 2017]. Proliferating LSCs have a turnover rate *p*_*Y*_ modulated by a logistic growth with carrying capacity *T*_*Y*_. They enter the quiescent compartment with inactivation rate *p*_*YX*_. Parameter *e*_*TKI*_ in describes the effect of TKI on proliferating LSCs. A linear dose-response effect is assumed, according to studies that verified such relationship within the range of usual imatinib treatment doses (100-800mg) [La Rosee et al., 2004, Deininger et al., 2003]. The TKI treatment is applied from treatment start at time *t* = 0 until treatment cessation at time *t* = *t*_*C*_; the function *u*(*t* − *t*_*C*_) is the unit-step function, *u*(*t* − *t*_*C*_) = 0 if *t* < *t*_*C*_ and *u*(*t* − *t*_*C*_) = 1 if *t* > *t*_*C*_. Immune cells are produced at a constant rate *p*_*Z*_ and are degraded at a natural rate *d*_*Z*_. In the absence of leukemic cells, immune cells reach the steady state *p*_*Z*_*/d*_*Z*_.

The interactions between leukemic and immune cells are described as follows. The immune effector response, i.e., the targeting of proliferating LSCs by immune cells, is described by *m*_*K*_*ZF*(*Y*). Here, *m*_*K*_ is the maximum rate at which one immune cell kills the leukemic cells, in units of (leukemic cells)/(immune cell × time). The function *F*(*Y*) is a non-dimensional quantity ranging between 0 and 1, and describes the modulation of the targeting of LSCs by the immune cells, as a function of the number of proliferating LSCs. The different choices for *F*(*Y*) are discussed below.

Conversely, the effect of LSCs on the stimulation and/or the inhibition of immune cells (summarized as recruitment) is described by the term *m*_*R*_*ZG*(*Y*). Following the mechanistic model derived by Kuznetsov et al. [1994], we assume that a stimulation occurs when one immune cell encounters leukemic cells and release signaling molecules to recruit more immune cells, while the inhibition occurs due to the releasing of anti-immune signals by LSCs. Parameter *m*_*R*_ represents the maximum recruitment rate at which one immune cell stimulates other immune cells, in units of 1/time. Alternatively, *m*_*R*_ can be understood in units of (immune cells)/(immune cell × time), thereby depicting the per capita recruitment by each immune cell. The function *G*(*Y*) is a non-dimensional quantity ranging between 0 and 1, and represents the modulation of the immune recruitment, as a function of the number of proliferating LSCs.

Our model assumes that leukemic stem cells do not directly target or compete with normal hamatopoietic cells. However, the model considers an implicit competition mechanism between normal and leukemic cells, by assuming that the sum of normal and leukemic cells is constant. Under this assumption each new leukemic cell replaces a normal cell (see [Fassoni et al., 2018] for more details). With this hypothesis, the model reflects the fact that, at CML diagnosis, almost only leukemic cells are found within the bone marrow, and the number of normal cells increases as the TKI-treatment proceeds with reduction of tumor load.

Finally, we assume that the quiescent leukemic cells are predominantly found in the bone marrow niches and are largely protected from the immune cells. For this reason, we also assume that the contribution of inactive leukemic cells to the stimulation of immune cells is negligible.

### 2.1 Modeling the immune response against leukemic cells

We consider four different functional responses for the function *F*(*Y*) modulating the targeting of leukemic cells by immune cells: A) a linear response, described by the law mass-action (or Holling type I); B) a response with saturation effect, described by the Holling type II functional response; C) a response with saturation and learning effects, described by the Holling type III functional response; and D) a response with an optimal immune window, recently suggested by Clapp and colleagues [Clapp et al., 2015]. Each of these choices represents different underlying biological mechanisms as shown in Figure 2.

**Figure 2:**
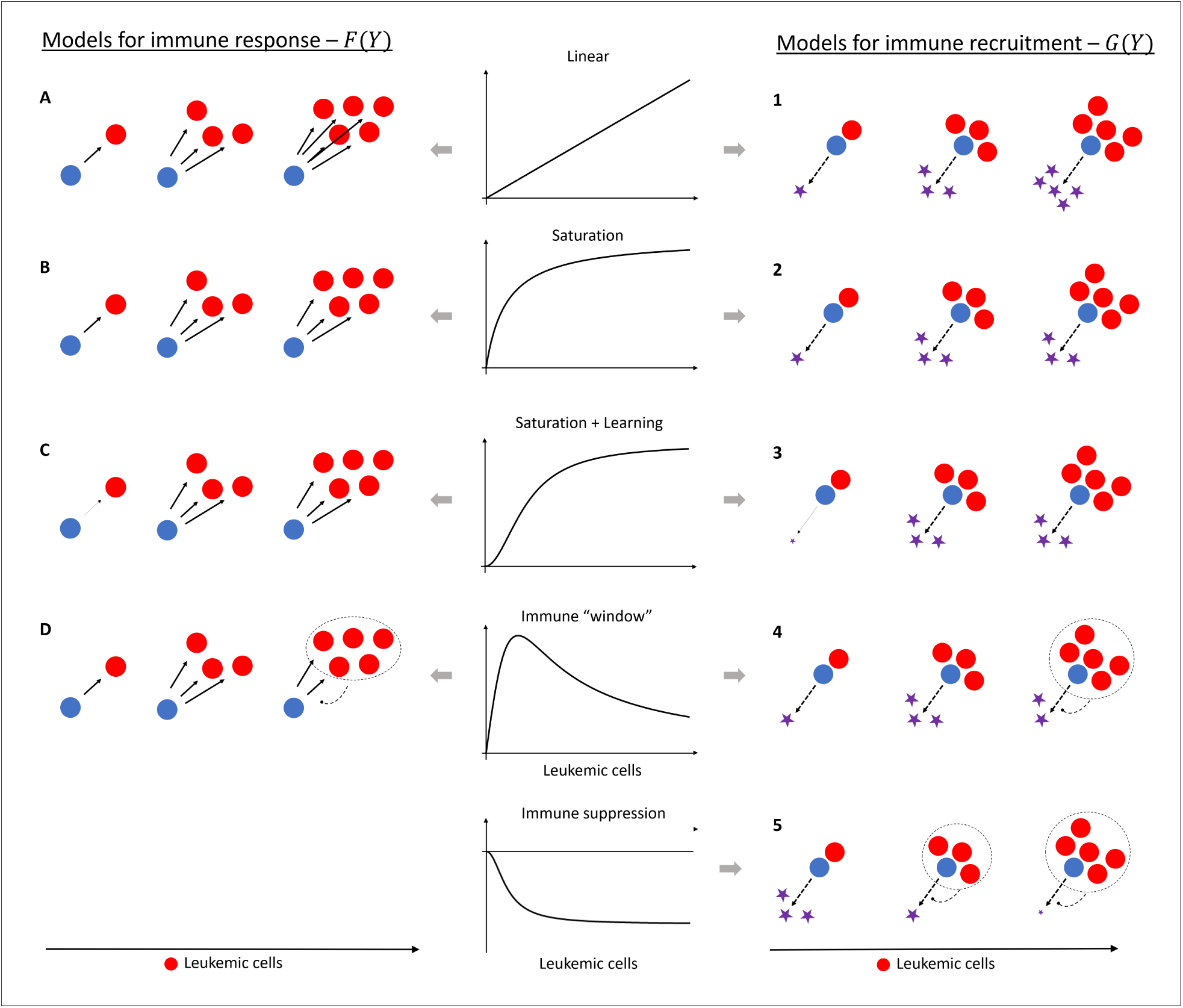
Schematic representation of the different functional responses describing the immune response and immune recruitment: the law of mass action, describing a linear response (first row); the Holling type II functional response, describing a saturation effect (second row); the Holling type III functional response, describing saturation and learning effects (third row); the functional response suggested by Clapp et al. [2015], describing an optimal immune window (fourth row); and a negative Holling type III functional response, describing a suppression of immune recruitment for high tumor load (fifth row).

#### 2.1.1 Law of mass action - Linear response

The law of mass action states that the removal of leukemic cells is linear with respect the number of LSCs (Figure 2, first row), i.e.,

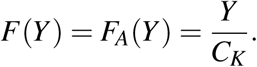

Here, the coefficient *C*_*K*_ represents the level of LSCs at which the immune attack reaches its maximum value, i.e., *F*(*C*_*K*_) = 1. This functional expression for the modulation of the immune attack implies that more leukemic cells lead to a higher efficiency of the targeting of a single immune cell.

#### 2.1.2 Holling type II - Saturation effect

The Holling type II functional response is a modification of the law of mass action. It assumes that the killing of leukemic cells is linear for low densities but, after a threshold described by the parameter *C*_*K*_, there is a saturation of the capacity of immune cells (Figure 2, second row). The expression for *F*(*Y*) in this case is

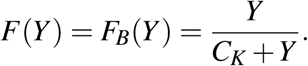

*C*_*K*_ corresponds to the number of LSCs at which the immune response reaches half of its capacity, i.e., *F*(*C*_*K*_) = 1*/*2. *F*(*Y*) saturates at 1 for high values of *Y*.

#### 2.1.3 Holling type III - Saturation and learning effects

The Holling type III functional response is similar to the Holling type II. The difference occurs for low densities, when the targeting of leukemic cells is very small (Figure 2, third row). In ecology, this functional response is used to describe learning effects in the predator ability, or also a mechanism of prey switching [Holling, 1959]. In our context, this choice indicates that, in the very beginning of leukemic growth, the immune system does not recognize the leukemic cells as a threat and does not target these cells. As the number of leukemic clones increases, there is an improvement of the immune response, representing effects such as acquired immunogenicity. The formal expression in this case is

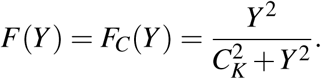

Again, the parameter *C*_*K*_ corresponds to the level of LSCs at which the response reach half of its maximum value, i.e., *F*(*C*_*K*_) = 1*/*2, and defines the threshold separating the learning and saturation effect.

#### 2.1.4 Immune window

This functional response has been used in a recent model study on CML [Clapp et al., 2015] and reflects the assumption that, although the immune response increases as the leukemic cell number increases, there is an inhibition of the immune system for a large population of leukemic cells (Figure 2, fourth row). As a result, we have a “window” for the number of leukemic cell at which the immune system works more efficiently. The expression for this term is

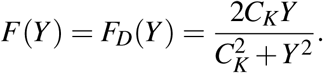

The parameter *C*_*K*_ controls the position where the maximum of *F*(*Y*) occurs. The factor 2*C*_*K*_ in the numerator is included for notational reasons, in order to keep *F*(*Y*) a non-dimensional quantity and to assure that the maximum is *F*(*C*_*K*_) = 1.

### 2.2 Modeling the recruitment of immune cells

Once an immune cell encounters and targets leukemic cells, it also promotes further immune cells to be activated and attack the tumor. The function *G*(*Y*) modulates the maximum number *m*_*R*_ of new immune cells recruited per time, per each immune cell in contact with *y* leukemic cells. To describe the possible mechanisms behind this recruitment, we consider the same structures as for the functions *F*(*Y*) above, and an additional response shape based on recent biological evidence pointing to the suppression of immune effector cell number by CML cells [Hughes et al., 2017] (see Figure 2).

#### 2.2.1 Law of mass action

The first possibility is to choose a linear recruitment

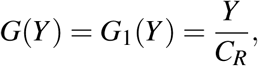

which means that the immune recruitment is proportional to the number of leukemic cells targeted by one immune cell (Figure 2, first row).

#### 2.2.2 Holling type II - Saturation effect

Choosing a Holling type II functional response,

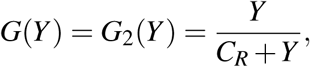

we have a saturation of the signaling capacity of immune cells (Figure 2, second row). This model for recruitment was derived by Kuznetsov et al. [1994] with reasonable assumptions on the mechanisms of release of signaling molecules by immune cells. The behavior described by this functional response reflects the assumption that there is a limited number of receptors on the surface of immune cells which sense the presence of leukemic cells, or there is a limited number of proteins in the signaling cascade which propagates the input signals inside each cell until the recruitment signaling is released in the extra-cellular medium.

#### 2.2.3 Holling type III – Saturation and learning effects

The choice of a Holling type III function for the recruitment,

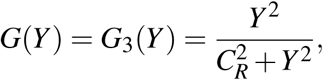

is a modification of the Holling type II, reflecting the assumption that the immune cells do not release almost any signal when a low numbers of leukemic cells are present and interactions are rare (Figure 2, third row). This choice reflects a type of adaptive response, in the sense that the immune system does not considers a low number of leukemic cells as an ‘emergency’ and does not trigger further activation unless the number of leukemic cells reaches a certain level.

#### 2.2.4 Immune window

The fourth possible function describing immune recruitment is given by

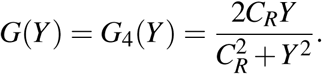

In the context of recruitment, the window obtained from this function reflects the assumption that a high number of leukemic cells contributes to the inhibition of immune recruitment (Figure 2, fourth row).

#### 2.2.5 Immune suppression

The last function used to describe modulation of the number of immune cells by the tumor load reflects recent biological findings pointing towards the immunosuppressive role of CML cells, exerted by different markers on the surface of CML cells which suppress the number of immune cells as the disease progresses [Hughes et al., 2017] (Figure 2, fifth row). This modulation can be described by

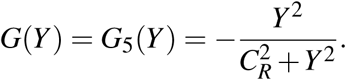

This choice assumes that the normal immune recruitment in the absence of CML cells, described by the natural influx *p*_*Z*_, is suppressed for increasing numbers of CML cells, resulting in a smaller net recruitment *p*_*Z*_ − *m*_*R*_*ZG*(*Y*). The parameter *C*_*R*_ controls the level of tumor load at which the suppression on the recruitment reaches half of its maximum absolute value, i.e., *G*_5_(*C*_*R*_) = *−*1*/*2.

## 3 Results

### 3.1 Quantitative criteria of “data fitting” are not sufficient for model selection

Combining all possible choices for immune response *F*(*Y*) and immune recruitment *G*(*Y*), we obtain 20 different sub-models. It is our primary objective to identify which of these models better represent the disease and treatment dynamics of CML. As a first criteria for model selection, we consider the ability of each model to fit individual patient time courses.

The tumor load in TKI-treated CML patients is usually monitored by the ratio BCR-ABL1/ABL1 in the peripheral blood, which measures BCR-ABL1 transcript levels by real time - PCR analysis. The relative easiness and robustness of these measurements makes CML a well monitored disease and suitable for quantitative modeling. To compare our model simulations with patient data, we assume that the ratio BCR-ABL1/ABL1 in the peripheral blood is equivalent to the percentage of proliferating LSCs in the stem-cell compartment. Thus, the modeled BCR-ABL1/ABL1 ratio, *L*_*MOD*_(*t*), which will be compared with the observed patient data, is given by

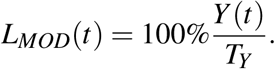

In order to illustrate the ability of each sub-model to optimally describe patient data, we took the time-course of one representative patient of the IRIS trial as an example, and applied parameter estimation methods to identify, for each of the 20 sub-models, the particular parameter values for which the model simulation *L*_*MOD*_(*t*) reproduces the observed data (Figure 3). The choice of parameter values and the fitting strategy are presented in Appendix B. We conclude that, with respect to the quantitative criteria of data fitting, none of the 20 sub-models can be excluded, nor can they be classified as more or less suitable.

**Figure 3:**
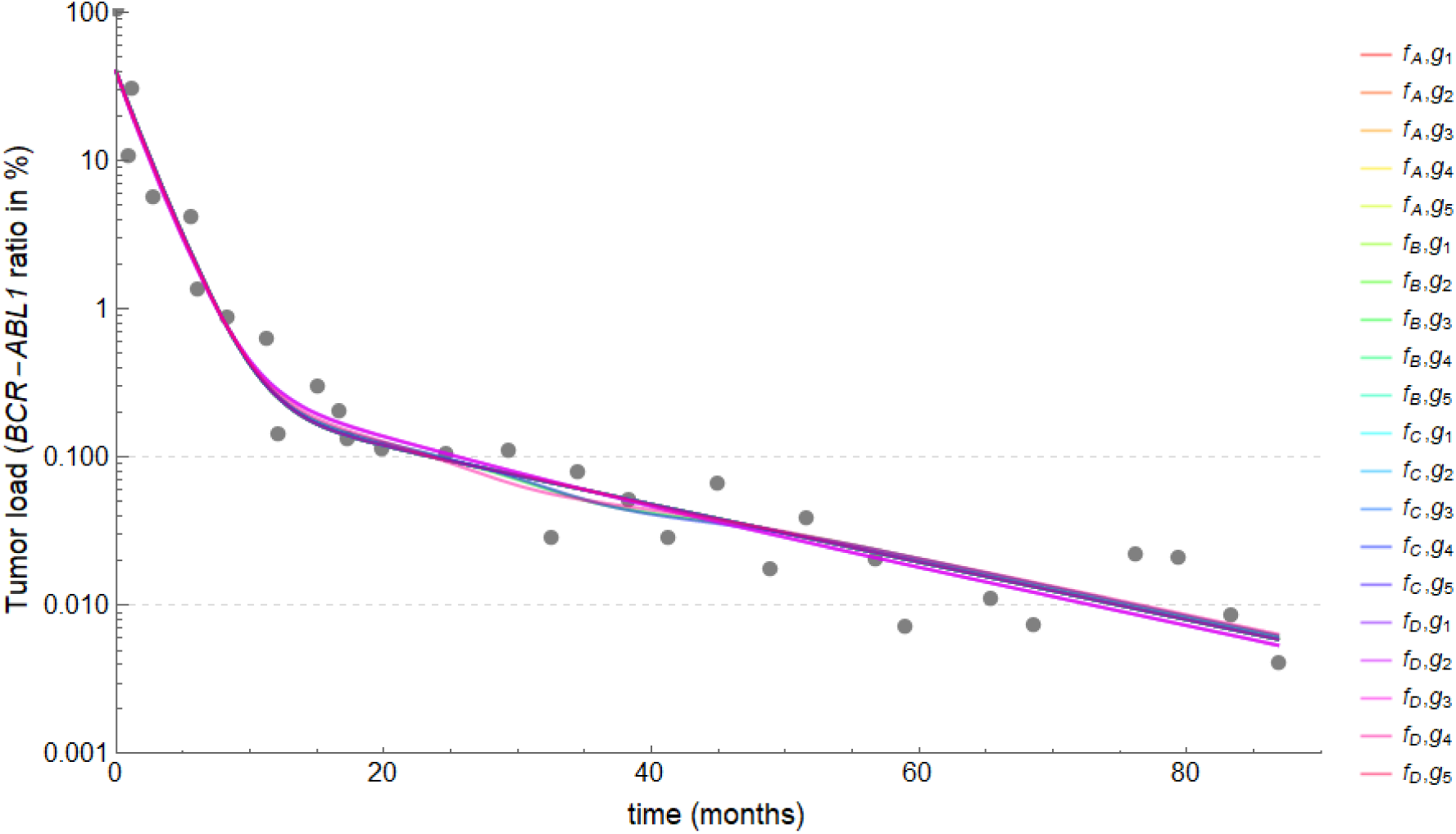
Fitting sub-models to patient data. Plots of the simulated BCR-ABL1/ABL1 ratio, *L*_*MOD*_(*t*), for each of the 20 sub-models (different colors), fitted the observed patient data (gray dots).

### 3.2 Qualitative criteria for model selection and a mathematical framework for model analysis

The above results imply that quantitative model fits are not sufficient to distinguish models with respect to their ability to explain time course data as all models perform sufficiently well (as in Figure 3). Instead we suggest including additional qualitative criteria to distinguish models based on their ability to account for general dynamic phenomena of disease patterning. Such dynamic phenomena are most obvious under system perturbations, such as changes in the treatment schedule, and are imprinted into the phase portrait of each sub-model. A complete mathematical analysis of each submodel is beyond the scope of this work, but it is our objective to investigate which of these models are suited to reflect the different phenomena observed in CML patients after treatment cessation. The results of this qualitative analysis will provide insights on the mechanisms underlying CML dynamics. The qualitative criteria are formalized as follows.

In different clinical trials to evaluate treatment cessation, about 40-50% of the eligible patients relapsed while the other 50-60% of patients remained in sustained remission [Saussele et al., 2016]. Among the later, some patients did not exhibit any detectable level of BCR-ABL1, suggesting that the disease might have been completely eradicated, while other patients exhibit very low, although detectable levels of BCR-ABL1, even many years after cessation. This proportion was around 30% in the A-STIM study [Rousselot et al., 2014]. It has been hypothesized that the immune system might play a major role in controlling or even eradicating these residual leukemic cell levels [Hughes and Yong, 2017].

From the dynamical systems point of view, such an immunological control suggests the existence of two or even three different attractors, corresponding to different CML levels: the “disease steady state” (corresponding to CML growth prior to diagnosis or to relapse), the “remission steady state” (corresponding to a sustained molecular remission) and potentially even a “disease-free steady state” (corresponding to cure or extremely low, thus undetectable, disease levels). In this context, the disease is diagnosed in the attracting region (or basin of attraction) of the disease steady state, while TKI treatment is depicted as a temporary force moving the system trajectories towards the attracting regions of the other steady states. If the treatment indeed succeeds to lead the system to one of these attracting regions (molecular remission or cure), the system will be maintained in such remission/cure region after treatment cessation. Otherwise, the system remains in the disease region and results in a relapse. From the clinical point of view, “cure” and “undetectable disease” might be the same state, due to incapability of measuring low disease levels accurately.

To analyze the ability of each model to qualitatively describe CML dynamics, we study the behavior and phase portrait (stability of steady states, basins of attraction and separatrices) of all submodels defined by equation (1) without treatment. The treatment-free system (i.e. *e*_*TKI*_ = 0 or *t* > *t*_*C*_) is denoted as the “permanent system” and describes the dynamics after treatment cessation. We will investigate which of the 20 sub-models present structurally different phase portraits that correspond to different clinically relevant scenarios for CML dynamics after TKI stopping. To describe these phase portraits and their corresponding clinical scenarios, we formally introduce the three types of steady states:

1. The “disease steady state” is a nontrivial stable equilibrium point for the permanent system, denoted as *E*_*H*_ = (*X*_*H*_, *Y*_*H*_, *Z*_*H*_), where *Y*_*H*_ corresponds to a high level of proliferating LSCs (*H* for high), near to their carrying capacity, i.e., 0 *≪ Y*_*H*_ ~ *T*_*Y*_.
2. The “remission steady state” is another nontrivial stable equilibrium point for the permanent system, denoted as *E*_*L*_ = (*X*_*L*_,*Y*_*L*_, *Z*_*L*_), where the number of proliferating LSCs is low (*L* for low), and corresponds to detectable BCR-ABL1 levels above background *Y*_*L*_ *≈* 10^*−*3^*T*_*Y*_ *≪ Y*_*H*_.
3. The “cure steady state” is the trivial equilibrium of the permanent system, given by *E*_0_ = (0, 0, *Z*_0_), and corresponds to the extinction of CML cells and the presence of immune cells at the normal level *Z*_0_ = *p*_*Z*_*/d*_*Z*_. In some situations, there is a third nontrivial steady state *E*_*D*_ = (*X*_*D*_,*Y*_*D*_, *Z*_*D*_) which is stable (together with *E*_*H*_ and *E*_*L*_, while *E*_0_ is unstable) and corresponds to a very low number of proliferating LSCs, with BCR-ABL1 levels below the detection limit, i.e., *Y*_*D*_ *≈* 10^*−*5^*T*_*Y*_ *" Y*_*L*_. We assume that it represents an undetectable residual level of BCR-ABL1 by current RT-PCR analysis, and will denote *E*_*D*_ as the “deep remission steady state” (*D* for deep) describing a “functional cure”.

When a positive nontrivial steady state is a saddle, it will be denoted by *E*_*S*_; the invariant manifolds of these saddle-points will determine the separatrix between the basins of attraction.

Based on the above notation, we define six structurally different phase portraits, which a submodel may exhibit. Each phase portrait corresponds to a unique clinically relevant scenario, which is expected to occur in the clinical situation. The scenarios are:

I. Scenario I - The phase portrait of the permanent system presents only two steady states: the disease steady state *E*_*H*_ is globally stable for positive solutions, and the cure steady state *E*_0_ is unstable. In this case, any solution of the permanent system with a nonzero initial number of proliferating LSCs (*Y* (0) > 0) will be attracted by the disease steady state *E*_*H*_. Therefore, a relapse is always expected after treatment cessation, because (mathematically) there is always a residual leukemic population after any treatment (Figure 4A).
II. Scenario II - The disease steady state *E*_*H*_ and the cure steady state *E*_0_ are the stable steady states in the permanent system. In this case, the treatment may result in a complete cure or a relapse after treatment cessation, depending on the duration of the treatment and the configuration of the basins of attraction of *E*_*H*_ and *E*_0_ (Figure 4B).
III. Scenario III - the disease steady state *E*_*H*_ and the remission steady state *E*_*L*_ are the stable steady states in the permanent system. In this case, remission or relapse after treatment cessation (but not a complete cure) can be described by the model, depending on the duration of the treatment and the configuration of the basins of attraction of *E*_*H*_ and *E*_*L*_ (Figure 4C).
IV. Scenario IV - The permanent system exhibits three stable steady states: the disease steady state *E*_*H*_, the remission steady state *E*_*L*_ and the cure steady state *E*_0_. In this case, treatment can lead to all possible outcomes, i.e., relapse, remission and cure, depending on the duration of the treatment and the configuration of the basins of attraction of *E*_*H*_, *E*_*L*_ and *E*_0_ (Figure 4D).
V. Scenario V: in this scenario, the cure steady state *E*_0_ is unstable, while other three nontrivial steady states are stable, i.e., the deep-remission steady state *E*_*D*_, the remission steady state *E*_*L*_ and the disease steady state *E*_*H*_ are stable. Clinically, this scenario might be indistinguishable from scenario IV, because this additional deep-remission steady state *E*_*D*_ can be interpreted as equivalent to the cure steady state *E*_0_ due to the very low number of CML cells (Figure 4E).

ø. Scenario ø - For completeness, we define an additional scenario where the cure steady state *E*_0_ is globally stable in the permanent system. This scenario corresponds to a healthy individual, where CML onset is impossible (Figure 4F).

**Figure 4:**
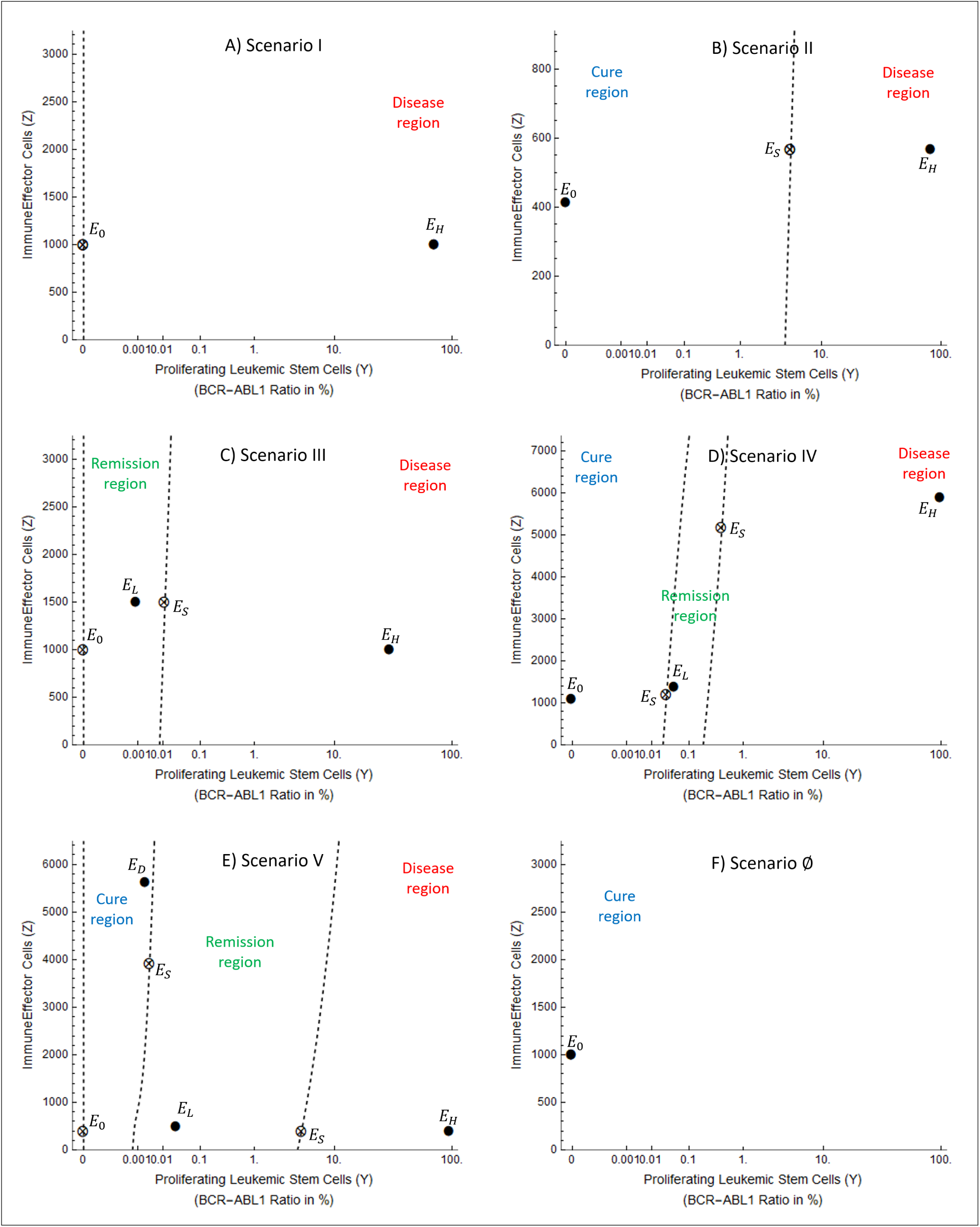
Possible clinically relevant scenarios (phase portraits). Representations of the three-dimensional phase space (*X,Y, Z*) for the permanent system (equation (1) with *e*_*TKI*_ = 0), restricted to the two-dimensional plane *X* = (*p*_*YX*_*/p*_*X*_*Y*)*Y*. Stable steady states are represented by solid circles (•) while saddle steady states are represented by crossed circles (⊗). The separatrix between the basins of attraction are represented by dashed lines. There is one basin of attraction in scenarios I and ø, two basins of attraction in scenarios II and III, and three basins of attraction in scenarios IV and V. Due to the huge difference in values of *Y* for the different steady states, and due to the impossibility to represent the value *Y* = 0 with the log-scale, the horizontal axis *Y* is set to a nonlinear scale via the transformation 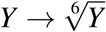.

With this nomenclature, the roman numerals in each scenario corresponds to the total number of positive nontrivial steady states (see Figure 4, e. g. scenario V corresponds to 5 nontrivial positive steady states: *E*_*H*_, *E*_*L*_, *E*_*D*_ and two saddle-points *E*_*S*_).

We analyzed each of the 20 sub-models with respect to the question, whether or not it can generate the clinically relevant scenarios for different parameter combinations. To do so, we adopted the following approach: for each sub-model, we mathematically analyzed the possible number of non-trivial steady states which could be exhibited. Since each of the five scenarios corresponds to a configuration with a characteristic number of non-trivial steady states (see Figure 4), we can mathematically rule out some of the scenarios for each sub-model. For instance, we have shown that the sub-model corresponding to choices (*F, G*) = (*F*_*C*_, *G*_5_) always has at least one nontrivial steady state, but it cannot exhibit exactly 2 or exactly 4 non-trivial steady states. This allows to conclude that this particular sub-model does not exhibit scenarios II and IV, and that the *possible* scenarios for this submodel are scenarios I, III and V. After this mathematical analysis, we numerically investigate whether such *possible* scenarios are actually exhibited by each sub-model. For the example above (sub-model (*F, G*) = (*F*_*C*_, *G*_5_)) we aimed to identify three different sets of biologically plausible parameter values that reproduce each of the possible scenarios (I, III and V) with respect the existence of the characteristic number of non-trivial steady states and their stability. In the end, we obtained a complete description of all scenarios actually exhibited by each sub-model. The results are summarized in Table 1. The detailed proofs and methods used this approach are described in Appendix A. As the next step we selected those sub-models that are suited to reproduce the clinical phenomena observed in CML patients.

**Table 1:** Scenarios (phase portraits) exhibited by each sub-model. Each combination of *F*(*Y*) = *F*_*i*_(*Y*), *i* ∈{*A, B,C, D*} and *G*(*Y*) = *G*_*j*_(*Y*), *j* ∈{1, 2, 3, 4, 5}, results in a structurally different sub-model of the permanent system described by system (1) with *e*_*TKI*_ = 0. This table presents the scenarios actually exhibited by each of the 20 sub-models, meaning that, for each sub-model (*F*_*i*_, *G*_*j*_) it is possible to choose different combinations of biologically plausible parameter values leading to the different indicated scenarios. See the description of each scenario in Figure 4. Sub-models selected according to the criterion in section 3.3 are marked by orange and green colors, which distinguish them into two classes: the *two-basins* (orange) and the *three-basins* (green) models (see section 3.4). The results in this table were obtained by using analytic and numerical methods presented in Appendix A.

| | $F_A$ | $F_B$ | $F_C$ | $F_D$ |
| --- | --- | --- | --- | --- |
| $G_1$ | $\emptyset, I$ | $\emptyset, I, II$ | $I, III$ | $\emptyset, I, II, III$ |
| $G_2$ | $\emptyset, I$ | $\emptyset, I, II$ | $I, III$ | $\emptyset, I, II, III$ |
| $G_3$ | $\emptyset, I$ | $\emptyset, I, II$ | $I, III$ | $\emptyset, I, II, III$ |
| $G_4$ | $\emptyset, I, III$ | $\emptyset, I, II, III, IV$ | $I, III, V$ | $\emptyset, I, II, III, IV, V$ |
| $G_5$ | $\emptyset, I, II, III$ | $\emptyset, I, II$ | $I, III, V$ | $\emptyset, I, II, III, IV$ |

### 3.3 Criterion for model selection: comparison with clinical outcomes after treatment cessation

The criterion which we impose for selecting a sub-model is whether it is able or not to reproduce the three clinical outcomes after treatment cessation, namely relapse, sustained remission and cure. In the following, we rule out the sub-models which do not exhibit these three possible outcomes.

First, we note that scenario ø is not a clinically relevant scenario. Although cure is achieved in this scenario, it is also achieved without treatment, because *E*_0_ is globally stable in the permanent system with *e*_*TKI*_ = 0. Thus, such scenario cannot represent a CML patient, as is does not describe CML diagnosis and progression in the absence of treatment.

As seen in Table 1, a linear killing effect (i.e., *F* = *F*_*A*_) combined with choices *G*_1_, *G*_2_, *G*_3_ and *G*_4_ does not reproduce scenarios where cure is possible (excluding scenario ø), because such sub-models reproduce only scenario I and III. On the other hand, the linear killing effect reproduces scenarios I, II and III when a suppression mechanism is chosen for the immune recruitment (*G* = *G*_5_) and reproduces the expected clinical outcomes (relapse, remission, cure). Therefore, we exclude the choices *F* = *F*_*A*_ with *G* ≠ *G*_5_ from the list of potential candidate mechanisms.

When a Holling type II functional response is chosen for the immune response to CML (*F* = *F*_*B*_) only one sub-model fits the criteria (model (*F*_*B*_, *G*_4_)). The other choices reproduce only scenarios I and II, and do not predict the clinical outcome of molecular remission. Therefore, these choices (*F* = *F*_*B*_ with *G* ≠ *G*_4_) are excluded from the list of potential candidates.

The choice of a learning effect in the immune response to CML (*F* = *F*_*C*_) leads to scenarios where cure is structurally impossible when *G* = *G*_1_, *G*_2_, *G*_3_, because such sub-models exhibit only scenarios I and III. On the other hand, the choices (*F*_*C*_, *G*_4_) and (*F*_*C*_, *G*_5_) additionally exhibit scenario V, which is viewed as a version of scenario IV where the cure-equilibrium is replaced by the deepremission-equilibrium *E*_*D*_. Therefore, by the first criteria, sub-models (*F*_*C*_, *G*_1_), (*F*_*C*_, *G*_2_) and (*F*_*C*_, *G*_3_) are excluded from the list of potential candidates, while the other remain.

Finally, when the “immune window” response (*F* = *F*_*D*_) is chosen, all the possible combinations for immune recruitment lead to sub-models that reproduce the clinically expected scenarios (relapse, remission, cure).

### 3.4 Models classification according to the need of bifurcations to describe clinical situations

After selecting the sub-models which in principle reproduce the possible clinical outcomes, we further distinguish them into two different classes. The first class corresponds to sub-models which are able reproduce the clinical outcomes with a static parameter set, while the second class comprises submodels which need to transit between different parameter sets to describe all the three possible clinical outcomes after treatment cessation, namely relapse, remission, or cure.

Scenarios IV or V correspond to phase portraits in which all clinically relevant situations (i.e. relapse, remission and cure) coexist for a particular parameter set. In such scenarios, two patients with identical model parameters could potentially end up in different “outcomes” if the treatment intensity or duration “drives” the trajectory into distinct basins of attraction. We defined the first class of sub-models as those models which exhibit at least one of scenarios IV or V. We refer to these sub-models as “three-basins models”; these are marked by a green color in Table 1.

In the other side, there are the sub-models which exhibit only scenarios I, II and III (marked by an orange color in Table 1) and in which either the remission, the cure, or none of these states is available for a particular parameter configuration. Here, one would need to use different parameter values to describe patients with different outcomes (cure or remission). This implies the need for bifurcations to change the qualitative phase portrait. We refer to these sub-models, which exhibit scenarios I, II and III only, as “two-basins models”.

Table I further indicates that all three-basins models also exhibit scenarios II and III. In this respect, they offer more complexity, but cannot rule out the occurrence of bifurcations during treatment. We do not argue in favor of three-basins or two-basins models. However, we point out the differences between the sub-models and reflect them in the light of certain biologically relevant question. Let’s consider the following question: Is every patient who reaches sustained treatment-free molecular remission a virtually curable patient, if she/he receives a more intensive therapy? If yes, we expect that a suitable CML model shall always present three stable steady states, favoring the three-basins models. If one argues that certain boundary conditions, like a patient’s particular immune system is generally not suited to allow a cure (even for a more intensive therapy), one may then suggest the two-basins models. In this case, a change of parameters (such as an additional sensitization of the immune system) may result in a bifurcation and, thereby, a qualitatively altered phase space, which then yields to another phase portrait in which the cure steady state is stable. Such a qualitative alteration of the phase space could possibly be supported by the administration of immune-modulatory drugs that act independent from the TKI.

### 3.5 Models classification according to different ways of reproducing the onset of CML

We now turn our attention to the pre-treatment phase of CML, i.e., the time interval from the onset of the first leukemic clone (i.e., mutation of a single cell) until diagnosis of a macroscopic disease. Based on the hypotheses and interactions considered by our general ODE model, we argue that it is plausible to use the outlined models to describe also this pre-treatment phase of CML.

To do so, we make the hypotheses that CML arises from a single leukemic stem cell initiated within a population of immune cells at a normal, unchallenged level. In the permanent system (equations (1) with *e*_*TKI*_ = 0), this is described by the initial conditions *X* (0) = 0, *Y* (0) = 1 cell and *Z*(0) = *p*_*Z*_*/d*_*Z*_. Therefore, the onset of CML is represented by a very small perturbation from the cure steady state *E*_0_. The persistence of the first clone and its progeny is only possible if *E*_0_ is unstable; otherwise, this small perturbation would return to *E*_0_, i.e., the immune system would control the onset of CML.

We showed that the cure steady state *E*_0_ is stable if, and only if, parameters satisfy

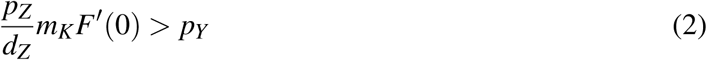

The proof is presented in Appendix A.3. The biological interpretation is the following: the value *m*_*K*_*F*^*t*^(0) measures the rate at which one immune cell mounts a response against the onset of the first leukemic cell. This value multiplied by *p*_*Z*_*/d*_*Z*_ measures the response rate of the entire immune population in the very beginning of leukemic growth. If this rate is greater than the proliferation rate of the leukemic cells, *p*_*Y*_, then the immune system is able to control the onset of leukemia. If not, the leukemic cells are able to break the first immune barrier and grow to a certain level. Interestingly, this condition does not depend on the recruitment function *G*(*Y*), but only on the immune cells already resident at the time of leukemic onset.

It is worth to note that the above condition cannot be satisfied when the immune response is given by *F*(*Y*) = *F*_*C*_(*Y*), since 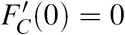. This choice represents an immune system which takes time to learn that the leukemic cells must be targeted. This delay, due to the learning effect, leads to an inability to control the onset of CML.

Now, to fully describe the onset and diagnosis, once we assume *E*_0_ is unstable and CML cells have escaped the first immune barrier, it is necessary that they grow until reaching the diagnosis size. This means that *E*_0_ must be in the attracting basin of *E*_*H*_, so that the trajectory starting at the small perturbation from *E*_0_ moves towards *E*_*H*_. Otherwise the solution would converge to another nontrivial steady state, such as *E*_*L*_ or *E*_*D*_, and the “disease progression” would not be observed. There are only two possible scenarios (phase portraits) which fit into this description and where the onset of CML is fully covered.

#### 3.5.1 Onset of CML starting from scenario I: the CML timeline viewed as sequential transitions through different phase-portraits

The first possibility is to start from scenario I, where *E*_*H*_ is globally attracting for positive solutions (Figure 5). In this case, the system needs to be pre-set into scenario I at the time of occurrence of this first leukemic episode. This means that, at this time and stage, the immune system is not able to control disease onset. However, if the phase portrait remains unchanged during disease progression, then, after diagnosis and treatment, a relapse would be always observable. As this contradicts clinical observations, we argue that there must be a change in the phase portrait during disease progression, i.e., a bifurcation from scenario I to another scenario must occur such that at the time of diagnosis the system has other, additional stable steady states. Possible transitions are from scenario I to II (via a transcritical bifurcation involving *E*_0_ and *E*_*L*_) or from scenario I to III (via a saddle-node bifurcation involving *E*_*L*_ and *E*_*U*_); see Figure 5. Further transitions may occur from scenario II to IV, from III to V (via saddle-node bifurcations) or between II to III ((via a transcritical bifurcation involving *E*_0_ and *E*_*L*_). All these bifurcations would be attributed to changes in parameters during disease progression. As a plausible example, we point out that the strength of the immunological response against CML may increase as the disease progresses, due to adaptative or stimulation effects on the immune system. This would be reflected by an increase in parameter *m*_*K*_. This increase may lead to the bifurcations pointed out in Figure 5, through scenarios I-II-IV or I-III-V, as exemplified in the bifurcation diagrams in Figure 6. However, assuming that the system converged to the disease steady state *E*_*H*_ before the bifurcation occured it will remain there as the immunological response is too slow and too late to change the trajectory towards the newly established basin of attraction for remission or cure.

**Figure 5:**
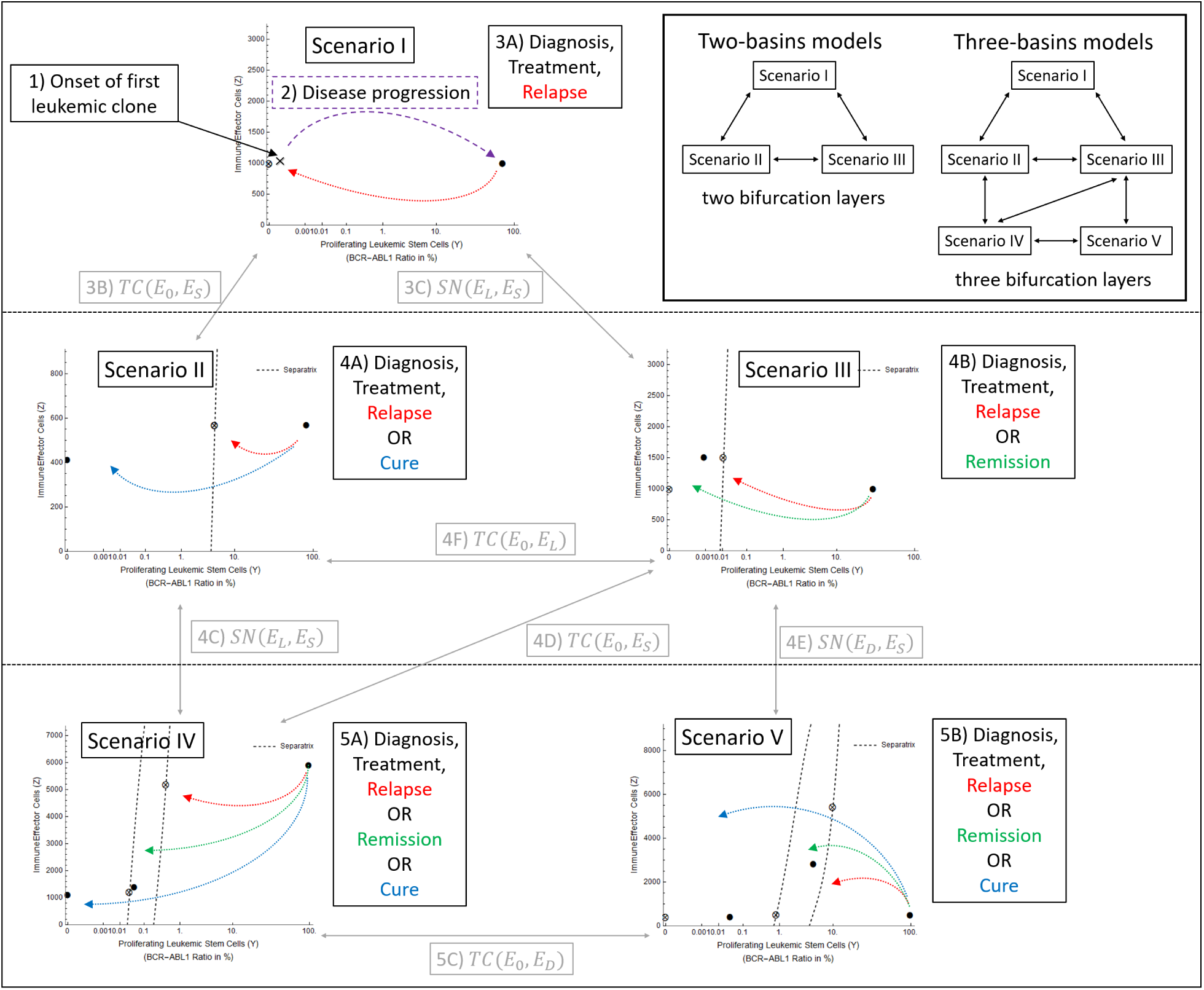
Routes from CML onset until diagnosis, treatment, and treatment cessation; starting from Scenario I. Some of the possible routes starting from scenario I and passing through scenarios II, III, IV and V, are: 1-2-3A (stays in scenario I), 1 → 2 → 3B → 4A (until scenario II; see Figure 6A), 1 → 2 → 3C → 4B (until scenario III; see Figure 6B), 1 → 2 → 3C → 4D → 5A (until scenario IV; see Figure 6C), and 1 → 2 → 3C → 4E → 5B (until scenario V; see Figure 6D). *TC*(*E*_0_, *E*_*J*_) denotes a transcritical bifurcation between steady states *E*_0_ and *E*_*J*_ (*J* = *S, D* or *L*), while *SN*(*E*_*J*_, *E*_*S*_) denotes a saddle-node bifurcation between *E*_*J*_ (*J* = *L* or *D*) and a saddle-point *E*_*S*_. Upper-right part: the two-basins models, which exhibit only scenarios I, II and III, transit along two bifurcation layers; while the three-basins models, which additionally exhibit scenarios IV or V, transit along three bifurcation layers.

**Figure 6:**
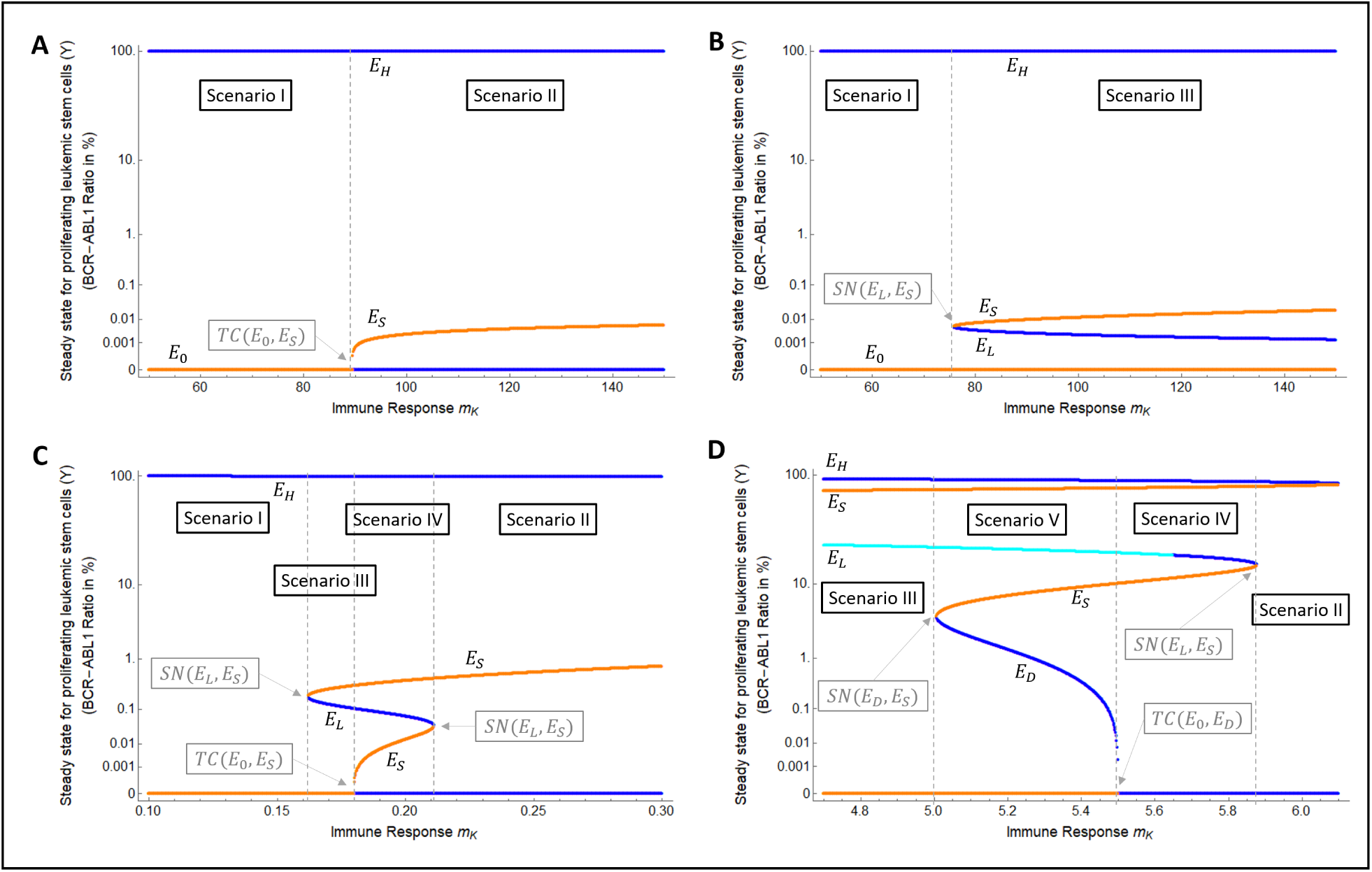
Numerical bifurcation diagrams illustrating changes in the attractor landscapes. Yellow curves correspond to saddle steady states, blue curves correspond to stable steady states (dark blue for stable nodes, light blue for stable focus). These bifurcation diagrams illustrate the different possibilities of transition of sub-models through different phase portraits as the immunological parameters change; the bifurcation points are indicated by vertical dashed lines. Each panel shows a different path as indicated in the caption of Figure 5. In all cases shown here, the bifurcation parameter is the strength of immune response *m*_*K*_, for which a small value of the leads to scenario I, where the onset of CML is possible. Similar transitions are obtained by varying the proliferation rate *p*_*Y*_ of LSCs and keeping *m*_*K*_ constant (not shown). A) I → II. B) I → III. C) I → III → IV → II. D) III → V → IV→ II. These bifurcation diagrams were obtained using an automated numerical routine which evaluates the steady states for each value of the bifurcation parameter. For each of this steady states, the eigenvalues of the corresponding jacobian matrix were calculated, and the steady states were classified according to the usual linear stability criteria.

As we stated before, the system needs to be in scenario I at the moment of CML onset in order to observe disease progression. However, it is not necessary that this scenario is the “natural” condition for every patient. Indeed, the system may be initially in other scenarios where disease initiation is not possible (when *E*_0_ is stable, as in scenarios II, IV or even ø), or not visible (when *E*_0_ is unstable but belongs to the attracting basin of a remission steady state, as in scenarios III and V). In such cases, a bifurcation leading to scenario I is required prior to onset and may be caused by a temporary loss or decrease in the immunological function, reducing parameter *m*_*K*_. Once this disruption is caused and the system moves to scenario I, the disease initiates, and may not be controlled anymore, even with a return to the original scenario due to a regain of immunological response, as commented above.

#### 3.5.2 Onset of CML starting from scenario III*: the complete CML timeline described by a single scenario

The second possibility to describe CML onset with model (1) is to consider an alternative phase portrait to scenario III, which we denote by III*. In this phase portrait, the stable steady states are *E*_*H*_ and *E*_*L*_, as is the case of scenario III, but now the cure steady state *E*_0_ is in the basin of attraction of *E*_*H*_ (Figure 7A). In this new scenario, a small perturbation from *E*_0_, such as the onset of one CML cell, will be attracted to *E*_*H*_, leading to disease progression and diagnosis. After treatment start, the system will be moved to the basin of attraction of *E*_*L*_, leading to remission (Figure 7A, blue dotted lines), or will remain in the basin of attraction of *E*_*H*_, leading to relapse (Figure 7A, red dotted lines). The actual outcome depends on the treatment parameters (intensity and duration). Therefore, besides reproducing the onset of CML until diagnosis, this scenario already reproduces the clinical outcomes of remission or relapse after treatment, without the need of a bifurcation to pass to another scenario.

**Figure 7:**
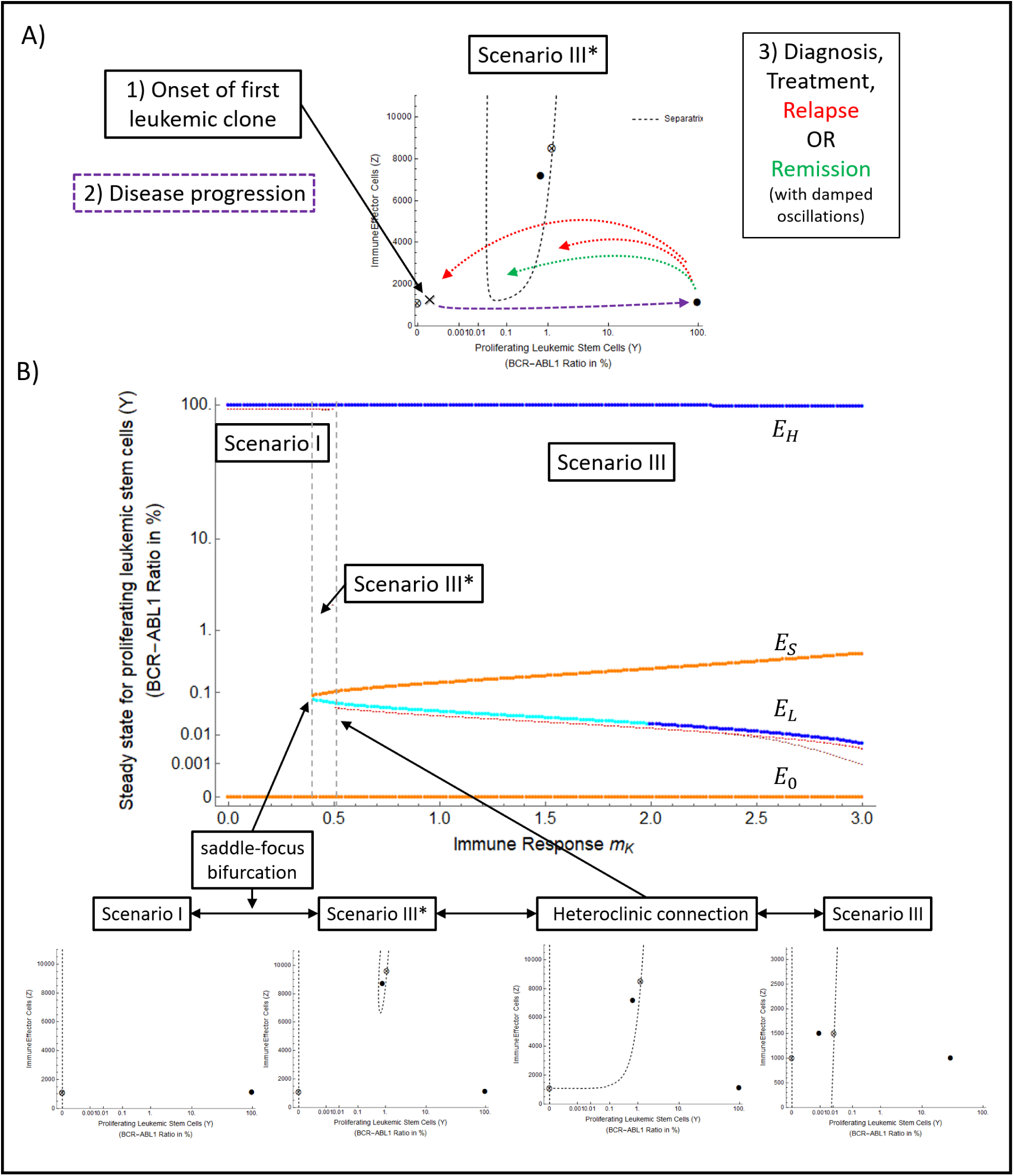
Onset of CML starting from scenario III*. (A) Route from CML onset until diagnosis, treatment, and treatment cessation. Scenario III* explains CML onset, progression, treatment and relapse/remission without the need of any bifurcation. (B) Formation of scenario III*. Scenario III* originates when the system is in scenario I and a parameter change (in this example, an increase in the immune response *m*_*K*_) leads to a saddle-focus bifurcation between *E*_*S*_ and *E*_*L*_. On the other hand, scenario III* also emerges when the system is in scenario III, and a parameter change (in this example, a decrease in the immune response *m*_*K*_) leads to a heteroclinic connection between the invariant manifolds of *E*_*S*_ and *E*_0_, so that the basin of attraction of *E*_*L*_ passes through a dramatic change in size and shape, and the cure steady state (unstable) *E*_0_ turns to belong to the basin of attraction of *E*_*L*_. In general, the parameter interval for which scenario III* is exhibited is narrow (it is interval between the two dashed lines), meaning that small changes in parameters could lead to scenarios I or III.

Interestingly, we observe that, in scenario III*, a very long or intense treatment may pass the basin of attraction of *E*_*L*_ and end near *E*_0_ but in the basin of attraction of *E*_*H*_, therefore, leading to relapse, whereas a weaker or shorter treatment would lead to remission (Figure 7A, red dashed line). Biologically, this corresponds to a situation where the immune system is capable to control the residual CML cells if these are not too few. In contrast, the immune system is tuned down if there is very low number of CML cells and, once they grow and activate the immune response again, it is too late to control them.

Also worth noting, the remission steady state *E*_*L*_ is a stable focus in scenario III*, which means that, once a successful treatment drives the system inside the basin of attraction of *E*_*L*_, the trajectories converge to it with damped oscillations. Although this fact was verified numerically, the shape of the basin of attraction of *E*_*L*_ within scenario III*, and the saddle-focus bifurcation involved in the arising of scenario III* are evidences suggesting that *E*_*L*_ is always a focus within this scenario. This implies that oscillating BCR-ABL1 dynamics should be observable during and after treatment, until the patient reaches TFR (Figure 6B). This oscillating phenomenon was reported during the treatment phase by Clapp et al. [2015]. Here, we show that these oscillations may be observable even after treatment cessation and may be a signature of an immune system which turns off and allows disease relapse when strong treatments eradicate too many CML cells. Indeed, in the model scenarios where TFR is reached without oscillations (scenarios II, III, IV or V), the cure steady state *E*_0_ does not belong to the basin of attraction of the disease steady state *E*_*H*_, and thus an additional time or dose in the treatment would not lead to relapse, contrary to scenario III*.

As a drawback of adopting scenario III* to describe the entire CML timeline, we point to the fact that this scenario does not predict a complete eradication of CML (*E*_0_ is always unstable). Therefore, patients in which no CML is detectable would, according to the model, still have a very low number of CML cells during the TFR phase. Furthermore, scenario III* is restricted to a narrow parameter interval, thereby strongly limiting its robustness (see Figure 7B). Additionally, there is no clinical evidence for an over-treatment in CML as the duration of TKI treatment has been repeatedly identified as a predictor of TFR. These limitations argue in favor of scenarios involving bifurcations to change the attractor landscape during disease progression and treatment.

## 4 Discussion

We presented a general mathematical model to describe the dynamics of CML with the particular focus on interactions with the immune response. Different treatment dynamics are described as trajectories within a landscape of possible steady states, which we refer to as disease states. Specifically, we consider 20 possible sub-models with different mechanistic assumptions on the interactions between CML and immune cells and show that a quantitative model adaptation to time course data before treatment cessation is not sufficient for model selection and needs to be complemented by the assessment of qualitative criteria. Introducing such criteria, we systematically compare the possible behaviors of each of the 20 sub-models with qualitative data on CML treatment and therapy cessation, and identified nine of them as more plausible compared to others. Under the hypothesis that our model is suitable to also describes CML onset and growth until diagnosis, we concluded that critical parameters of the interaction dynamics need to change during leukemia growth and treatment, thereby resulting in bifurcations that alter the attractor landscape of available and biologically meaningful disease states.

Within the nine sub-models satisfying the qualitative criteria, five assume the existence of a functional window for the immune response (function *f*_*D*_). The other four sub-models assume one of the other proposed functional forms of the immune response, but necessarily assume a recruitment function with either an immune window effect (function *g*_4_, 2 sub-models) or direct suppression of immune recruitment (function *g*_5_, 2 sub-models). Understanding that the immune window assumption describes an inhibited recruitment for high tumor load, we note that all the nine selected sub-models assume a suppression mechanism at least in one interaction (i.e., immune function or recruitment). This result agrees with recent data on immunological profiles of CML patients during TKI therapy [Hughes et al., 2017].

Indeed, several immune effector mechanisms appear to be inhibited or suppressed in CML patients at diagnosis, and are restored or enhanced during TKI treatment and after reaching deep molecular remission [Hughes and Yong, 2017]. Together with the immunosuppressive features of CML cells, these findings underline the functional dependencies of immune effector cell number and function, which we formally describe by the functional responses in the different selected sub-models. A further characterization of such functional responses, to be included in a future mathematical model, may be achieved by assessment of longitudinal data on the immune system of CML patients during continuing TKI treatment [Hughes et al., 2017].

To qualitatively assess the predictive power of our model, we simulated treatment cessation in all the nine selected sub-models. For each sub-model, we found two sets of parameters values, which reasonably well describe the data, but predict different outcomes: while one set leads to relapse after cessation (dashed lines in Figure 8), the other set leads to remission or cure (continuous lines in Figure 8). We obtained these two distinct outcomes by slightly varying the parameters *m*_*K*_ (maximum effector function of immune cells) and *p*_*Y*_ (leukemia proliferation rate) in each parameter set. Both parameters are crucial for the outcome of treatment cessation, since *m*_*k*_ directly affects the ability of immune cells to control the residual disease and *p*_*Y*_ determines the intrinsic aggressiveness of the leukemic cells. These results show that these parameters are not uniquely identifiable by the measurements of tumor load in response to TKI treatment only. Thus, additional measurements of immune cells function and number are necessary to correctly estimate these crucial parameters and to derive reliable model predictions. Therefore, besides informing the actual functional dependence between the immune effector cell number or function and the leukemic load, incorporating in mathematical models the longitudinal data on the immune system of CML patients could also make these models able to reliable estimate the outcome of treatment cessation in a patient-specific way.

**Figure 8:**
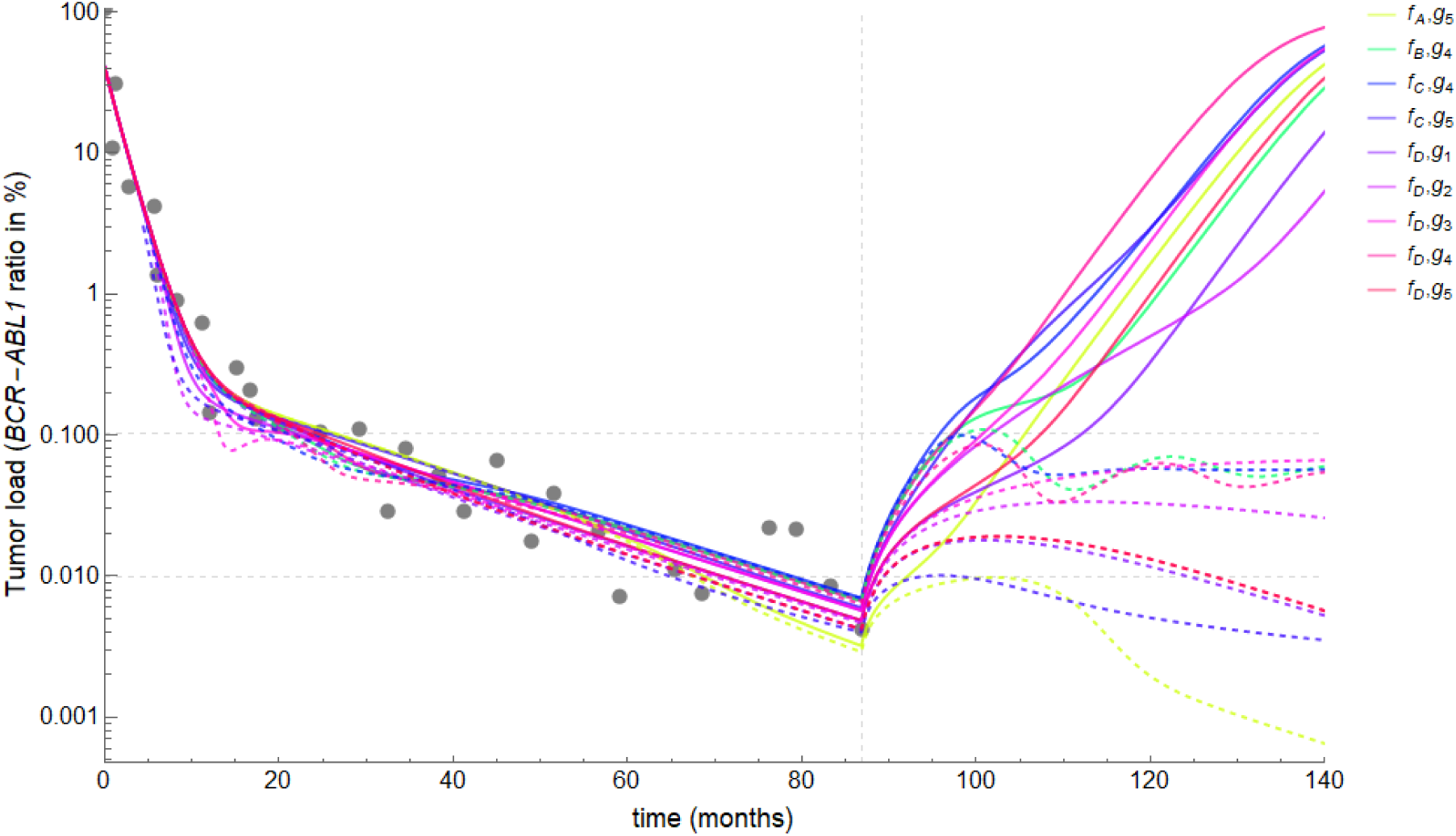
Unidentifiability of treatment response after theray cessation. Using model simulations to extrapolate patient data and predict outcome of treatment cessation. Plots of the modeled BCR-ABL1/ABL1 ratio, *L*_*MOD*_(*t*), for each of the 9 sub-models (different colors). For each sub-model, we identified two sets of parameter values which adjust the observed patient data (gray dots). When treatment cessation is simulated at the time of the last measurement, one parameter set leads to relapse (continuous lines) while the other leads to sustained remission/cure (dashed lines). Therefore, the crucial parameters for determining the outcome of treatment cessation are not identifiable with tumor load data only.

An alternative treatment strategy includes TKI-dose reduction by 50 % before treatment cessation, which has been recently studied the DESTINY trial [Clark et al., 2017]. Additionally, we discussed the potential of TKI dose reduction in the context of long-term maintenance treatment [Fassoni et al., 2018]. In both cases, the dose reduction is expected to cause a perturbation on the dynamics of CML, which can be captured and quantified by the patient-specific response in the BCR-ABL1/ABL1 ratios following dose reduction. This response may be informative about the patient’s immune system, and can provide an alternative or complementary information to be incorporated by mathematical models of CML -immune interactions.

Additionally, comparing the different phase portraits exhibited by the selected sub-models, we concluded that the full dynamics of pre-treatment, treatment and post-treatment phase can only be sufficiently and robustly covered if one assumes that critical model parameters change over time. Mathematically such parameter changes lead to bifurcations and introduce qualitative changes in the underlying attractor landscape. Our mathematical formalization allows to speculate about clinically relevant parameters that appear suitable to purposely deliver those changes. As such, the maximal intensity of immune effector cell activity *m*_*K*_ or the growth rate *p*_*Y*_ (quantifying leukemia aggressiveness) act as bifurcation parameters and can account for changes in the attractor landsca

Translating those findings into a biologically meaningful context, we suggest that if certain parameters, such as sensitivity of the immunological cells towards CML cells, can be identified as a target that changes during therapy, drug-based approaches (such as immune-modulatory drugs) may be implemented to alter the landscape of available disease states such that the remission or cure attractor emerges and increases in size. In this way, a TKI driven remission can be supported and permanently maintained even after therapy cessation. Earlier observations about the beneficial role of IFN-alpha treatment point towards this direction [Mahon et al., 2010].

Due to the homogeneity of the disease and the availability of high quality time course data, CML has attracted mathematical modeling approaches with different foci [Catlin et al., 2005, Michor et al., 2005, Roeder et al., 2006, Dingli et al., 2010, Stein et al., 2011, Komarova and Wodarz, 2009, Woywod et al., 2017]. Several models also studied the role of the immune system in CML and made particular assumptions about the underlying mechanisms [Kim et al., 2008, Wodarz, 2010, Clapp et al., 2015]. In our hands, there is no conclusive data to particularly highlight one over the other functional mechanisms of CML-immune interaction. For this reason we chose a broader approach and illustrate how a set of qualitative criteria is suited to rule out a set of very simple interaction mechanisms and points towards necessity of mutual suppression mechanisms between immune effector cells and leukemic cells. The model considered here has some simplifying assumptions, which allow to derive a mathematically analyzable model. First, the leukemic cells in the peripheral blood are not explicitly considered, but are assumed to be in number proportional to the number of proliferating LSCs [Stein et al., 2011, Fassoni et al., 2018]. Second, our ODE model does not consider stochastic effects, which might become relevant when the number of LSCs is very small and potentially lead to disease eradication even when the cure steady state is mathematically unstable. However, as such low LSCs numbers are only reached after very long treatment periods, we are confident to adhere to a regimen in which stochastic effects do not confer substantially different results. Finally, our model does not consider direct effects of the TKI itself on the immune system. Although there is some evidence for such an interaction [Zitvogel et al., 2016], it appears to be a second order effect in comparison with the effect on the immune response caused the TKI-induced reduction of tumor load [Hughes and Yong, 2017].

Our approach demonstrates that formalization in terms of mathematical models provide relevant complementary means to systematically study regulatory mechanisms and speculate about the role of immunological interactions in long term CML treatment. Our approach illustrates that the complexity of clinically observed phenotypes can only be explained if suppressive feedback mechanisms and dynamically altered interaction parameters are assumed. Experimental validation of those critical parameters opens the door to target them using appropriate drugs and to achieve sustained remission even after therapy cessation. Furthermore, mathematical models are an ideal tool for the virtual testing of new measurement protocols and treatment schedules, and can lead to the formulation of testable hypotheses.

## A Mathematical analysis

The general dynamics of system (1) depend on the interactions between leukemic and immune cells: four options for *F*(*Y*) and five options for *G*(*Y*) account for a total of 20 different combinations (sub-models). In this Appendix, we identify which scenarios (phase portraits ø, I, II, III, IV and V) are exhibited by each combination. Understanding that the TKI treatment acts as a temporary force moving the system trajectories in the phase space, we analyze the asymptotic behavior of the permanent system, i.e., system (1) with *e*_*TKI*_ = 0:

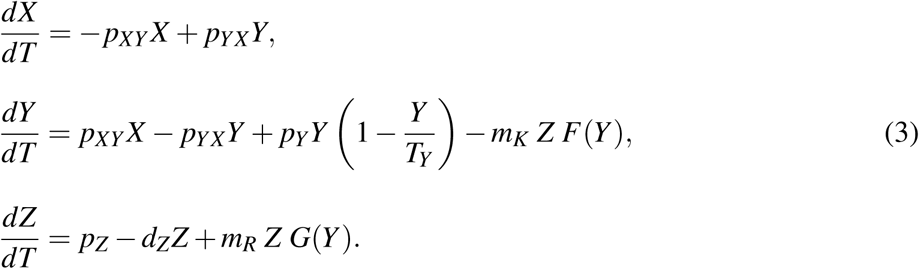

### A.1 Non-dimensionalization

To reduce the number of parameters in the analytic study, we introduce the non-dimensional variables

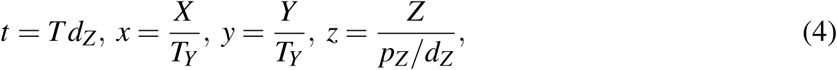

and obtain a non-dimensional model, mathematically equivalent to (3), given by

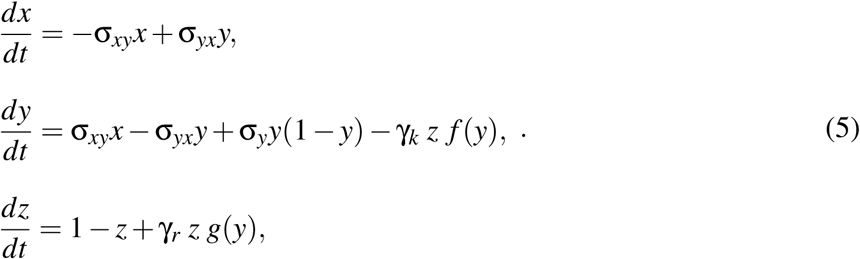

where the non-dimensional parameters are

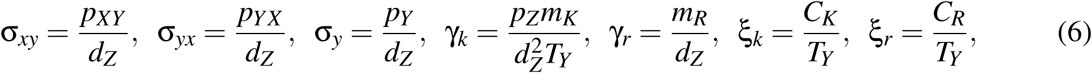

and the non-dimensional functional responses *f* (*y*) and *g*(*y*) are given, in each case *f*_*i*_(*y*) = *F*_*i*_(*T*_*y*_*y*), *i* ∈ {*A, B,C, D*} and *g* _*j*_(*y*) = *G*_*j*_(*T*_*y*_*y*), *j* ∈ {1, 2, 3, 4, 5}, by

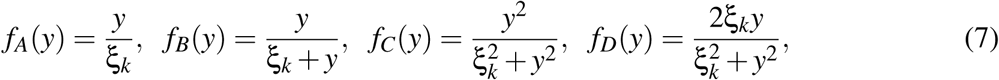

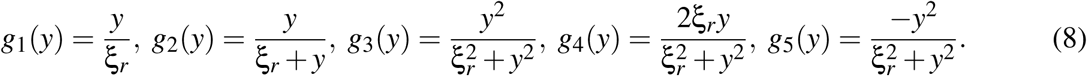

### A.2 Conditions on model parameters

We may impose biologically plausible conditions on some model parameters. As stated in Section 2, parameters *C*_*K*_ and *C*_*R*_ define the levels of leukemic cells at which the functions *F*_*i*_(*Y*) and *G*_*j*_(*Y*) exert their action. Thus, it is reasonable to assume that all these effects occur on a population level lower than the carrying capacity *T*_*Y*_ of the leukemic cells. Thus, we assume that *C*_*K*_,*C*_*R*_ < *T*_*Y*_. Further, we note that the expression 1*/d*_*Z*_ corresponds to the mean life-time of immune effector cells, while 1*/m*_*R*_ can be understood as the mean time of individual immune recruitment when the recruitment is at maximum, i.e., *G*(*Y*) ≈ 1. In other words, 1*/m*_*R*_ is the mean time spent by one immune cell between the last contact with leukemic cells and the arrival of a new immune cell, in a situation where the recruitment is at maximum. It is reasonable to assume that the former time (1*/d*_*Z*_) is greater than the latter (1*/m*_*R*_), i.e, *d*_*Z*_ < *m*_*R*_. Taken together, these conditions on the dimensional parameters, correspond to conditions on the non-dimensional parameters:

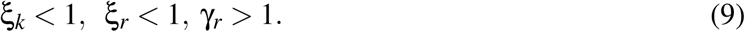

### A.3 Analysis of the trivial steady state *E*_0_

All models for immune response *f* (*y*) and recruitment *g*(*y*) satisfy *f* (0) = *g*(0) = 0. Thus, the point *E*_0_ = (0, 0, 1) (with coordinates (*x, y, z*)) is the trivial steady state for system (5). *E*_0_ is the “cure steady-state”. The Jacobian matrix of system (5) evaluated at *E*_0_ is

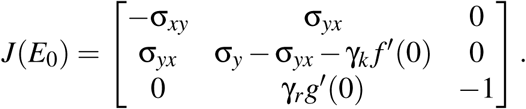

The characteristic polynomial of *J*(*E*_0_) is *p*_0_(λ) = *−*(λ + 1)(λ^2^ + *a*_1_λ + *a*_0_), where *a*_1_ = σ_*xy*_ + σ_*yx*_ + γ_*k*_ *f* ′ (0) *−* σ_*y*_ and *a*_0_ = σ_*xy*_ (γ_*k*_ *f* ′ (0) *−* σ_*y*_). Thus, one eigenvalue of *J*(*E*_0_) is λ_1_ = *−*1 < 0, while the other two satisfy λ_2_ + λ_3_ = *−a*_1_ and λ_2_λ_3_ = *a*_0_. We have the following cases:

1. If γ_*k*_ *f* ′ (0) > σ_*y*_ then *a*_0_ > 0 and *a*_1_ > 0. Thus, λ_2_, λ_3_ < 0 and *E*_0_ is a stable node.
2. If γ_*k*_ *f* ′ (0) < σ_*y*_ then *a*_0_ < 0. Thus, λ_2_ < 0 < λ_3_ and *E*_0_ is a saddle-point, with two negative eigenvalues.

The above results are summarized in the following proposition.

**Proposition 1.** *Let*

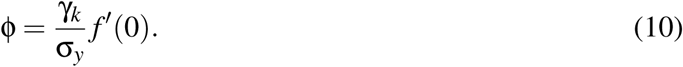

*If* ϕ > 1, *then E*_0_ *is a locally asymptotically stable steady state for system* (5). *If* ϕ < 1, *then E*_0_ *is a saddle-point for system* (5), *with one positive eigenvalue and two eigenvalues with negative real part.*

**Remark 1.** *In terms of the dimensional parameters, parameter* ϕ *is written as*

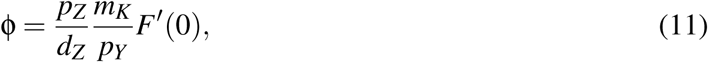

*(note that f* ′ (0) = *F*′ (0)*T*_*Y*_ *). Thus, the cure-equilibrium E*_0_ *is stable if, and only if*,

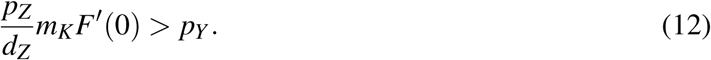

### A.4 Analysis of nontrivial steady states

Besides the trivial steady state, system (5) can have multiple other nontrivial steady states, which we study in the following. Setting *dx/dt* = 0 we obtain *x* = (σ_*yx*_*/*σ_*xy*_)*y*. Substituting this expression in *dy/dt* = 0 and solving for *z*, we obtain

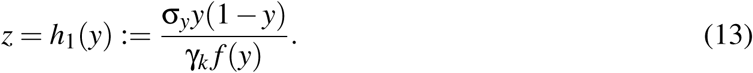

On the other hand, setting *dz/dt* = 0 leads to

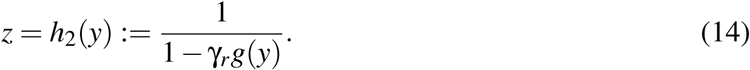

Taken together, (13) and (14) imply that the coordinate *y* ≠ 0 of a nontrivial steady state should satisfy

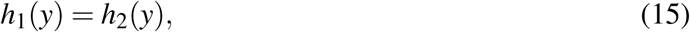

Therefore, the nontrivial steady states of system (5) are the points *E* = (*x*\*(*y*), *y, z*\*(*y*)) where

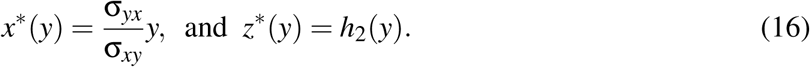

and *y* is a nonzero root of (15), which will be refereed as the equation for the nontrivial steady states. Further, a nontrivial steady state is biologically feasible if its coordinates are positive, i.e., *y* > 0, *x*\*(*y*) > 0 and *z*\*(*y*) ≥ 0. Checking expressions in (13) and (14) and using the fact that *f* (*y*) ≥ 0 for all choices of *f* (*y*), we conclude that a root *y* of *h*_1_(*y*) = *h*_2_(*y*) is feasible if, and only if, 0 < *y* ≤ 1 and *h*_2_(*y*) > 0. The results of this subsection are summarized in the following proposition.

**Proposition 2.** *The feasible nontrivial steady states of system* (5) *are the equilibrium points E* = (*x*\*(*y*), *y, z*\*(*y*)) *where x*\*(*y*) *and z*\*(*y*) *are defined in* (16) *and y is a root of* (15) *such that* 0 < *y* ≤ 1 *and h*_2_(*y*) > 0.

It is our goal to describe which qualitatively distinct phase portraits (classified as 5 different scenarios, defined in Figure 4) are exhibited by each sub-model and which are not. This question may be approached by assessing the number of roots of (15) which satisfy the feasibility conditions (note that scenario I corresponds to 1 feasible nontrivial steady state, scenario II corresponds to 2 feasible nontrivial steady states, and so on; see Figure 4). With this, we will be able to i) conclude that some scenarios are not exhibited by a given sub-model, due to the number of feasible nontrivial roots admitted by equation (15), and ii) conclude that some scenarios are possibly exhibited by such submodel. In the latter case, we will numerically confirm this possibility by obtaining parameter values for which the possible scenarios are indeed exhibited by the sub-model. Such analysis is summarized in the following theorem.

**Theorem 1.** *Consider system* (5) *under conditions* (9), *and scenarios* ø, *I, II, III, IV and V, defined by their number of feasible nontrivial steady-states, 0, 1, 2, 3, 4 and 5, respectively, with their local stability as indicated in Figure 4. Then, for each choice* (*f*, *g*) = (*f*_*i*_, *g* _*j*_), *i* ∈{*A, B,C, D* }, *j* ∈{1, 2, 3, 4, 5 }, *given in* (7) *and* (8), *a given subset of these scenarios cannot be exhibited by such sub-model, while the remaining scenarios do. Such subsets and the parameter values which provide the feasible scenarios are indicated in Tables 2 and 3.*

**Table 2:** Summary of analysis of models *F* = *F*_*A*_, *F*_*B*_ and *G* = *G*_1_, *G*_2_, *G*_3_, *G*_4_, *G*_5_. For each sub-model and each of scenarios ø, I, II, III, IV and V, we either present a set of parameter values which leads to the specified scenario, or exclude such scenario with a formal proof, presented in section A.5. See Appendix B for a reference to the other parameter values, which remained constant through all sub-models and scenarios.

| $F$ | $G$ | Scenario | $p_Y$ | $C_K$ | $C_R$ | $m_K$ | $m_R$ |
| --- | --- | --- | --- | --- | --- | --- | --- |
| $F_A$ | $G_1$ | ∅ | 0.2 | $5 \times 10^5$ | $5 \times 10^5$ | 110 | 2 |
| $F_A$ | $G_1$ | I | 0.2 | $5 \times 10^5$ | $5 \times 10^5$ | 10 | 2 |
| $F_A$ | $G_1$ | II, III, IV, V excluded by analytic tools | | | | | |
| $F_A$ | $G_2$ | ∅ | 0.2 | $5 \times 10^5$ | $5 \times 10^5$ | 110 | 2 |
| $F_A$ | $G_2$ | I | 0.2 | $5 \times 10^5$ | $5 \times 10^5$ | 10 | 2 |
| $F_A$ | $G_2$ | II, III, IV, V excluded by analytic tools | | | | | |
| $F_A$ | $G_3$ | ∅ | 0.2 | $5 \times 10^5$ | $10^4$ | 110 | 2 |
| $F_A$ | $G_3$ | I | 0.2 | $5 \times 10^5$ | $10^4$ | 10 | 2 |
| $F_A$ | $G_3$ | II, III, IV, V excluded by analytic tools | | | | | |
| $F_A$ | $G_4$ | ∅ | 0.2 | $5 \times 10^5$ | $10^3$ | 110 | 2 |
| $F_A$ | $G_4$ | I | 0.2 | $5 \times 10^5$ | $10^5$ | 90 | 2 |
| $F_A$ | $G_4$ | III | 0.2 | $5 \times 10^5$ | $10^5$ | 10 | 2 |
| $F_A$ | $G_4$ | II, IV, V excluded by analytic tools | | | | | |
| $F_A$ | $G_5$ | ∅ | 0.2 | $5 \times 10^5$ | $5 \times 10^5$ | 110 | 2 |
| $F_A$ | $G_5$ | I | 0.2 | $5 \times 10^5$ | $10^3$ | 90 | 2 |
| $F_A$ | $G_5$ | II | 0.2 | $5 \times 10^5$ | $10^3$ | 110 | 2 |
| $F_A$ | $G_5$ | III | 0.2 | $5 \times 10^5$ | $4 \times 10^5$ | 97 | 1.15 |
| $F_A$ | $G_5$ | IV, V excluded by analytic tools | | | | | |
| $F_B$ | $G_1$ | ∅ | 0.2 | $10^4$ | $2 \times 10^5$ | 6 | 2 |
| $F_B$ | $G_1$ | I | 0.2 | $10^4$ | $2 \times 10^5$ | 1.98 | 3 |
| $F_B$ | $G_1$ | II | 0.2 | $10^4$ | $5 \times 10^5$ | 4 | 2 |
| $F_B$ | $G_1$ | III, IV, V excluded by analytic tools | | | | | |
| $F_B$ | $G_2$ | ∅ | 0.2 | $10^4$ | $2 \times 10^5$ | 6 | 3 |
| $F_B$ | $G_2$ | I | 0.2 | $10^4$ | $2 \times 10^5$ | 1.98 | 4 |
| $F_B$ | $G_2$ | II | 0.2 | $10^4$ | $5 \times 10^5$ | 4 | 2 |
| $F_B$ | $G_2$ | III, IV, V excluded by analytic tools | | | | | |
| $F_B$ | $G_3$ | ∅ | 0.2 | $10^4$ | $2 \times 10^5$ | 6 | 12 |
| $F_B$ | $G_3$ | I | 0.2 | $10^4$ | $2 \times 10^5$ | 1.98 | 12 |
| $F_B$ | $G_3$ | II | 0.2 | $10^4$ | $5 \times 10^5$ | 4 | 2 |
| $F_B$ | $G_3$ | III, IV, V excluded by analytic tools | | | | | |
| $F_B$ | $G_4$ | ∅ | 0.2 | $10^4$ | $2 \times 10^5$ | 6 | 12 |
| $F_B$ | $G_4$ | I | 0.2 | $10^4$ | $2 \times 10^5$ | 1.98 | 12 |
| $F_B$ | $G_4$ | II | 0.2 | $10^4$ | $5 \times 10^5$ | 4 | 2 |
| $F_B$ | $G_4$ | III | 0.2 | $10^4$ | $10^5$ | 1.9 | 2 |
| $F_B$ | $G_4$ | IV | 0.2 | $10^4$ | $10^5$ | 2.1 | 2 |
| $F_B$ | $G_4$ | V excluded by analytic tools | | | | | |
| $F_B$ | $G_5$ | ∅ | 0.2 | $10^4$ | $10^3$ | 160 | 2 |
| $F_B$ | $G_5$ | I | 0.2 | $10^4$ | $10^3$ | 1.9 | 2 |
| $F_B$ | $G_5$ | II | 0.2 | $10^4$ | $10^3$ | 2.2 | 2 |
| $F_B$ | $G_5$ | III, IV, V excluded by analytic tools | | | | | |

**Table 3:** Summary of analysis of models *F* = *F*_*C*_, *F*_*D*_ and *G* = *G*_1_, *G*_2_, *G*_3_, *G*_4_, *G*_5_. For each sub-model and each of scenarios ø, I, II, III, IV and V, we either present a set of parameter values which leads to the specified scenario, or exclude such scenario with a formal proof, presented in section A.5. See Appendix B for a reference to the other parameter values, which remained constant through all sub-models and scenarios.

| $F$ | $G$ | Scenario | $p_Y$ | $C_K$ | $C_R$ | $m_K$ | $m_R$ |
| --- | --- | --- | --- | --- | --- | --- | --- |
| $F_C$ | $G_1$ | I | 0.2 | $5 \times 10^4$ | $10^5$ | 5 | 1.1 |
| $F_C$ | $G_1$ | III | 0.2 | $2 \times 10^4$ | $5 \times 10^5$ | 8 | 1.1 |
| $F_C$ | $G_1$ | $\emptyset, II, IV, V$ excluded by analytic tools | | | | | |
| $F_C$ | $G_2$ | I | 0.2 | $10^5$ | $10^5$ | 20 | 1.1 |
| $F_C$ | $G_2$ | III | 0.2 | $2 \times 10^4$ | $10^5$ | 8 | 1.1 |
| $F_C$ | $G_2$ | $\emptyset, II, IV, V$ excluded by analytic tools | | | | | |
| $F_C$ | $G_3$ | I | 0.2 | $10^5$ | $10^5$ | 20 | 1.1 |
| $F_C$ | $G_3$ | III | 0.2 | $2 \times 10^4$ | $10^5$ | 8 | 1.1 |
| $F_C$ | $G_3$ | $\emptyset, II, IV, V$ excluded by analytic tools | | | | | |
| $F_C$ | $G_4$ | I | 0.2 | $10^5$ | $2 \times 10^5$ | 20 | 1.5 |
| $F_C$ | $G_4$ | III | 0.2 | $10^5$ | $10^5$ | 20 | 1.1 |
| $F_C$ | $G_4$ | V | 0.2 | $1.55 \times 10^4$ | $1.7 \times 10^5$ | 4.805 | 1.01 |
| $F_C$ | $G_4$ | $\emptyset, II, IV$ excluded by analytic tools | | | | | |
| $F_C$ | $G_5$ | I | 0.2 | $10^5$ | $2 \times 10^5$ | 20 | 1.5 |
| $F_C$ | $G_5$ | III | 0.1 | $10^5$ | $10^5$ | 33.33 | 1.1 |
| $F_C$ | $G_5$ | V | 1 | $9.8 \times 10^4$ | $3.4 \times 10^4$ | 1800 | 10 |
| $F_C$ | $G_5$ | $\emptyset, II, IV$ excluded by analytic tools | | | | | |
| $F_D$ | $G_1$ | ∅ | 0.1 | $10^5$ | $4 \times 10^5$ | 10 | 1.1 |
| $F_D$ | $G_1$ | I | 0.1 | $10^5$ | $10^4$ | 0.5 | 1.1 |
| $F_D$ | $G_1$ | II | 0.1 | $10^5$ | $5 \times 10^5$ | 10 | 1.1 |
| $F_D$ | $G_1$ | III | 0.1 | $3 \times 10^4$ | $9 \times 10^4$ | 1.485 | 1.1 |
| $F_D$ | $G_1$ | IV, V excluded by analytic tools | | | | | |
| $F_D$ | $G_2$ | ∅ | 0.1 | $10^5$ | $10^5$ | 10 | 1.1 |
| $F_D$ | $G_2$ | I | 0.1 | $10^5$ | $10^4$ | 0.5 | 1.1 |
| $F_D$ | $G_2$ | II | 0.1 | $10^5$ | $5 \times 10^5$ | 10 | 1.1 |
| $F_D$ | $G_2$ | III | 0.1 | $3 \times 10^4$ | $9 \times 10^4$ | 1.485 | 1.1 |
| $F_D$ | $G_2$ | IV, V excluded by analytic tools | | | | | |
| $F_D$ | $G_3$ | ∅ | 0.1 | $10^5$ | $10^5$ | 10 | 1.1 |
| $F_D$ | $G_3$ | I | 0.1 | $10^5$ | $10^4$ | 0.5 | 1.1 |
| $F_D$ | $G_3$ | II | 0.1 | $10^5$ | $5 \times 10^5$ | 10 | 1.1 |
| $F_D$ | $G_3$ | III | 0.1 | $2 \times 10^5$ | $2.9 \times 10^5$ | 9.9 | 1.1 |
| $F_D$ | $G_3$ | IV, V excluded by analytic tools | | | | | |
| $F_D$ | $G_4$ | ∅ | 0.1 | $10^5$ | $5 \times 10^5$ | 10 | 1.1 |
| $F_D$ | $G_4$ | I | 0.1 | $2 \times 10^5$ | $5 \times 10^5$ | 9 | 1.1 |
| $F_D$ | $G_4$ | II | 0.1 | $10^5$ | $10^5$ | 10 | 1.1 |
| $F_D$ | $G_4$ | III | 0.1 | $10^5$ | $10^4$ | 0.5 | 1.1 |
| $F_D$ | $G_4$ | IV | 0.1 | $1.1 \times 10^5$ | $5 \times 10^5$ | 5.555 | 1.01 |
| $F_D$ | $G_4$ | V | 0.1 | $1.1 \times 10^5$ | $5 \times 10^5$ | 5.445 | 1.01 |
| $F_D$ | $G_5$ | ∅ | 0.5 | $1.3 \times 10^5$ | $10^5$ | 650 | 1.2 |
| $F_D$ | $G_5$ | I | 0.1 | $2 \times 10^5$ | $10^3$ | 9.9 | 1.01 |
| $F_D$ | $G_5$ | II | 0.1 | $10^5$ | $10^5$ | 10 | 1.1 |
| $F_D$ | $G_5$ | III | 0.1 | $3 \times 10^5$ | $5 \times 10^5$ | 14.85 | 1.1 |
| $F_D$ | $G_5$ | IV | 0.9 | $4.5 \times 10^5$ | $5 \times 10^2$ | 2800 | 13 |
| $F_D$ | $G_5$ | V excluded by analytic tools | | | | | |

**Remark 2.** *Theorem 1 is a ‘non-existence result’ which exclude the possibility of some sub-models to exhibit some scenarios. The affirmative answer is obtained by setting some model parameters (parameters p*_*Y*_, *C*_*K*_, *C*_*R*_, *m*_*K*_ *and m*_*R*_*) to a certain set of values in such way the scenario is observed when we numerically calculate the steady states and their local stability (through the numerical calculation of the jacobian matrix and its eigenvalues). The global behavior of system* (5) *(existence of limit cycles or other non-trivial attractors) is not approached by theorem 1. Table 1 summarizes the results of Tables 2 and 3.*

### A.5 Proof of theorem 1

To prove theorem 1, we first present the expressions of *h*_1_ and *h*_2_ for each choice of *f* (*y*) and *g*(*y*). We will express *h*_1_(*y*) in terms of ϕ, as this parameter is a threshold for the stability of *E*_0_ and so is related to changes between scenarios. Remembering that ϕ = (γ_*k*_*/*σ_*y*_) *f* ^*t*^(0) and using the formulas for *f*_*i*_(*y*), *i* ∈ {*A, B,C, D*} in equation (7), we obtain:

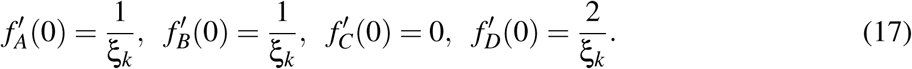

Then, ϕ = γ_*k*_*/*(σ_*y*_ξ_*k*_) if *f* = *f*_*A*_, ϕ = γ_*k*_*/*(σ_*y*_ξ_*k*_) if *f* = *f*_*B*_, ϕ = 0 if *f* = *f*_*C*_, and ϕ = 2γ_*k*_*/*(σ_*y*_ξ_*k*_) if *f* = *f*_*D*_. Thus, in each case, the formula for *h*_1_(*y*) can be expressed in terms of ϕ as

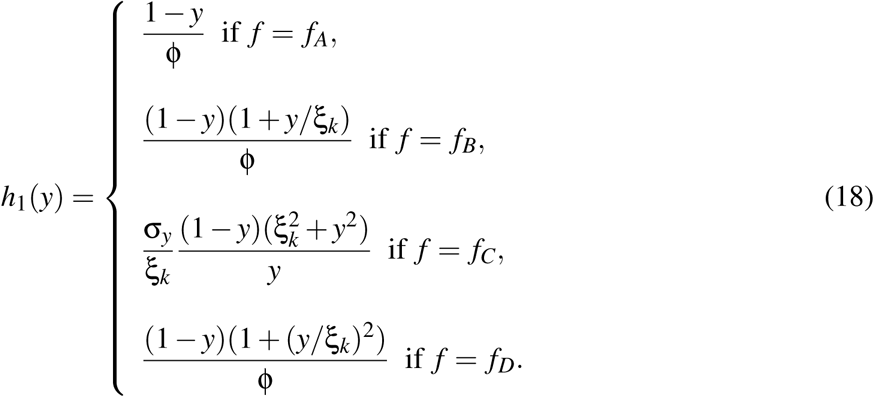

In the same way, the expressions for *h*_2_(*y*) in each case *g*(*y*) = *g* _*j*_(*y*), *j* ∈ {1, 2, 3, 4, 5}, are

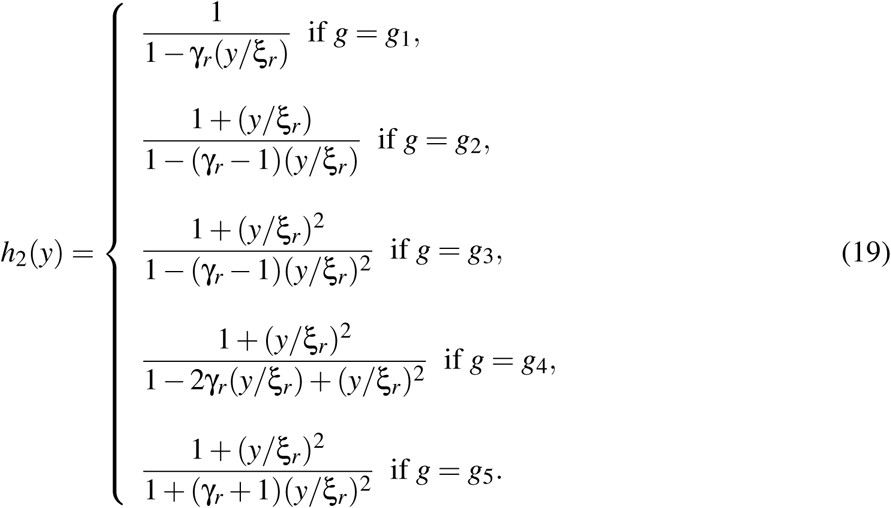

To analyze some of the sub-models, we will also study a polynomial equation equivalent to *h*_1_ = *h*_2_. From equations (18) and (19), we note that *h*_1_ and *h*_2_ are rational functions, i.e., quotient of polynomial functions. Therefore, equation *h*_1_(*y*) = *h*_2_(*y*), is equivalent to the polynomial equation

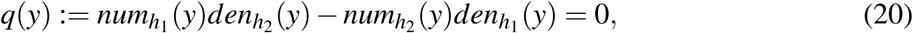

where 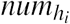 and 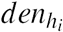 are the numerator and denominator of *h*_*i*_, respectively, and are polynomial functions. For each choice (*f*_*i*_, *g* _*j*_) of the sub-models, a different polynomial equation will be obtained, and the number of positive roots of such equation will be studied by using the *Descarte’s Rule of Sign*. If these roots satisfy the feasibility conditions, 0 < *y* ≤ 1 and *h*_2_(*y*) > 0, then, their quantity correspond to the number of feasible nontrivial steady states. The *Descarte’s Rule of Sign* states that the number of positive roots of a polynomial equation is equal to the number of sign variations between consecutive non-zero coefficients, or less than it by some even number [Kennedy, 2001]. For instance, if *q* is a fourth degree polynomial and its coefficients satisfy *c*_0_ > 0, *c*_1_ > 0, *c*_2_ < 0, *c*_3_ > 0 and *c*_4_ < 0, then the sign sequence is of *q* is *S*_*q*_ = (+ + − + −) (*S* for sequence) and the number of sign variations is *V*_*q*_ = 3 (*V* for variation). Therefore, such polynomial has 1 or 3 positive roots. The notation introduced in this paragraph will be used throughout the subsequent analysis. Further, we denote by ± the sign of a coefficient which can be positive or negative. To facilitate the referencing, we summarize the above results in the following lemma.

**Lemma 1.** *Let q*(*y*) = 0 *the polynomial equation for the nontrivial roots, with coefficients c*_*i*_. *The number of roots of q satisfying y* > 0 *is given by V*_*q*_ *−* 2*l, where l is a positive integer such that V*_*q*_ ≥ 2*l.*

Now, we introduce a way to easily verify which positive roots of *q* satisfy the feasibility condition *y* ≤ 1. Substituting *y* = 1 + *u* in *q*(*y*) = 0, we obtain another polynomial equation, *r*(*u*) = 0, where *r*(*u*) = *q*(1 + *u*). Each root *u* > 0 of *r*(*u*) = 0 corresponds to a root *y* = 1 + *u* > 1 of *q*(*y*) = 0. Thus, using the *Descarte’s Rule of Sign* as above, we may count the number of positive roots of *r*(*u*), which is equal to the number of positive roots of *q* which do not satisfy the feasibility condition *y* ≤ 1. Further, we note that *r*(*u*) has the same degree as *q* and, if *q*(*y*) = *c*_0_ + *c*_1_*y* + … + *c*_*n*_*y*^*n*^ and *r*(*u*) = *d*_0_ + *d*_1_*u* + … + *d*_*n*_*u*^*n*^ have degree *n*, then the coefficients *d*_0_ and *d*_*n*_ of *r*(*u*) are related to the coefficients *c*_*i*_ of *q* as

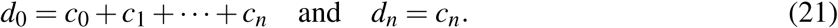

Furthermore, we conclude that, if *d*_0_ and *d*_*n*_ have different signs, then the sequence *S*_*r*_ of signs of *r*(*u*) has at least one sign variation and the number of sign variations *V*_*r*_ is an odd number. This is true because every sign change between *d*_0_ and *d*_*n*_ needs to occur in pairs and *V*_*r*_ = 1 + 2*k* for some integer *k* ≥ 0, i.e., *V*_*r*_ is an odd number if *d*_0_ and *d*_*n*_ have different signs. Using this result, we obtain a simple criterion for the existence of positive but non-feasible roots for *q*, which is summarized in the following *lemma*.

**Lemma 2.** *Let q*(*y*) = 0 *the polynomial equation for the nontrivial roots, with coefficients c*_*i*_, *and r*(*u*) = *q*(1 + *u*), *with coefficients d*_*i*_. *Then, the formulas in* (21) *hold. Furthermore, if d*_0_ *and d*_*n*_ *have different signs, then the number of the positive roots of q which satisfy the condition y* > 1 *is a positive odd number, less or equal the degree of q.*

We now analyze each sub-model in detail. Conditions (9) are assumed to hold.

**Analysis for** *f* = *f*_*A*_ **and** *g* = *g*_1_, *g*_2_, *g*_3_. For *f* (*y*) = *f*_*A*_(*y*), from (18), we have that

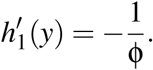

Therefore, *h*_1_(*y*) is a decreasing function. Furthermore, *h*_1_(*y*) is non-negative only in the interval 0 ≤ *y* < 1 with *h*_1_(0) = 1*/*ϕ and *h*_1_(1) = 0.

Now, we study *h*_2_(*y*) for the choices *g* = *g*_1_, *g*_2_, *g*_3_. From the expressions of *h*_2_(*y*) in these cases, we conclude the denominator of *h*_2_(*y*) vanishes and change its sig<u>n at an u</u>nique *y*_*d*_ > 0, where *y*_*d*_ = ξ_*r*_*/*γ_*r*_ for *g* = *g*_1_, *y*_*d*_ = ξ_*r*_*/*(γ_*r*_ − 1) > 0 for *g* = *g*_2_, and 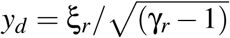 for *g* = *g*_3_ (since γ_*r*_ > 1). Analyzing the sign of the denominator, we conclude that *h*_2_(*y*) is positive for 0 ≤ *y* < *y*_*d*_ and is negative for *y* > *y*_*d*_. Therefore, the feasibility condition *h*_2_(*y*) > 0 restricts the analysis to the interval 0 ≤ *y* < *y*_*d*_. Now we show that *h*_2_(*y*) is increasing in this interval. From the general equation for *h*_2_(*y*) (14), we have

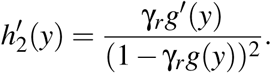

Since *g*(*y*) is an increasing function when *g* = *g*_1_, *g*_2_, *g*_3_, we have *g*′ (*y*) ≥ 0. Thus, 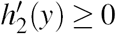 and *h*_2_(*y*) is an increasing function for these cases. Summarizing, in the interval 0 ≤ *y* < *y*_*d*_, we have *h*_2_(0) = 1, *h*_2_(*y*) is increasing, and *h*_2_(*y*) → +∞ when *y* → *y*_*d*_, *y* < *y*_*d*_.

Combining the properties of *h*_1_ and *h*_2_ we conclude the following. If *h*_1_(0) > *h*_2_(0), then *h*_1_ and *h*_2_ intersect in the interval 0 < *y* < *y*_*d*_ exactly once, at some *y*^*^ such that 0 < *y*^*^ < 1 and *h*_2_(*y*\*) > 0. Thus, there is an unique nontrivial steady state *E*. If *h*_1_(0) < *h*_2_(0) then *h*_1_ and *h*_2_ do not intersect and there is no feasible nontrivial steady state. Since *h*_1_(0) = 1*/*ϕ and *h*_2_(0) = 1, we have established the following result: if ϕ > 1, system (5) has an unique nontrivial steady state *E*; and If ϕ < 1, system (5) does not admit nontrivial steady states. This result states that sub-models with *f* = *f*_*A*_ and *g* = *g*_1_, *g*_2_, *g*_3_ do not exhibit scenarios II, III, IV and V. The only possible scenarios exhibited by these sub-models are I and ø. We numerically verified that such scenarios are indeed exhibited by these sub-models (see Table 2).

**Analysis for** *f* = *f*_*A*_ **and** *g* = *g*_4_. For *f* = *f*_*A*_, we have *h*_1_(*y*) = (1 − *y*)*/*ϕ. Thus, *h*_1_(*y*) is a decreasing function with *h*_1_(0) = 1*/*ϕ > *h*_1_(*y*) > *h*_1_(1) = 0 for 0 < *y* < 1. On the other hand, since *g*_4_(*y*) ≥ 0 *h*_2_(*y*) = 1*/*(1 − γ_*r*_*g*_4_(*y*)), we conclude that *h*_2_(*y*) ≥ 1 if *g*_4_(*y*) < 1*/*γ_*r*_ and, *h*_2_(*y*) < 0 if *g*_4_(*y*) > 1*/*γ_*r*_. Since one of the feasibility conditions is *h*_2_(*y*) > 0, we conclude that if a feasible root *y*^*^ occurs, then *h*_2_(*y*\*) ≥ 1. Hence, no feasible root occurs for ϕ > 1, since, in this case, 1 > *h*_1_(*y*) > 0 for 0 < *y* < 1. Therefore, only scenario ø occurs if ϕ > 1. The case ϕ < 1 is studied with the polynomial equation *q*, which has degree ∂(*q*) = 3 and coefficients

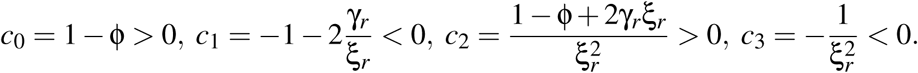

Thus, the sign sequence of *q* is *S*_*q*_ = (+ − + −) and the number of sign variations is *V*_*q*_ = 3. Therefore, *q* admits 1 or 3 positive roots. Thus, scenarios I and III are possible for model (*f*_*A*_, *g*_4_) if ϕ < 1, while the other scenarios are excluded. We numerically verified the occurrence of scenarios I and III (see Table 2).

**Analysis for** *f* = *f*_*A*_ **and** *g* = *g*_5_. In this case, *q* is a third degree polynomial with coefficients

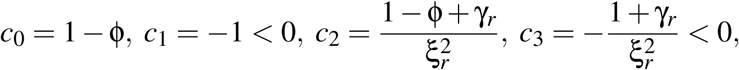

(remember that the nondimensional parameters are positive and satisfy conditions (9)). If ϕ < 1, the sign sequence is *S*_*q*_ = (+ − + −) and the number of sign variations is *V*_*q*_ = 3. Thus, such equation admits 1 or 3 positive roots and scenarios I and III are possible for model (*f*_*A*_, *g*_5_). If ϕ > 1, the sign sequence is *S*_*q*_ = (− − ± −) and the number of sign variations is *V*_*q*_ = 0 or 2. If we additionally have ϕ < 1 + γ_*r*_, then *c*_2_ > 0 and *V*_*q*_ = 2. In this case, equation *q*(*y*) = 0 admits 0 or 2 positive roots and scenarios ø and II are possible for model (*f*_*A*_, *g*_5_). We numerically verified the occurrence of scenarios I and III, and ø and II, under the respective conditions above (see Table 2).

**Analysis for** *f* = *f*_*B*_ **and** *g* = *g*_1_. In this case, *q* is a third degree polynomial with coefficients

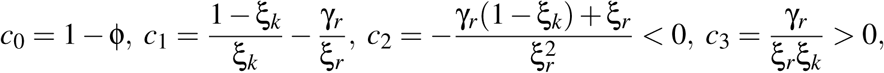

(remember that ξ_*k*_ < 1). By *lemma 2*, we obtain that the coefficients *d*_0_ and *d*_3_ of *r*(*u*) are

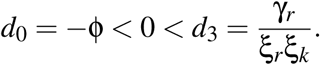

Thus, it follows also by *lemma 2* that 1 or 3 positive roots of *q* are non-feasible, i.e., satisfy *y* > 1. Now we analyze the sign sequence of *q*. If ϕ < 1, the sign sequence is *S*_*q*_ = (+ ± − +) and the number of sign variations is *V*_*q*_ = 2. Thus, such equation admits 0 or 2 positive roots. Since at least one is non-feasible, equation *q* has 2 positive roots but an unique feasible root. Thus, only scenario I is possible if ϕ < 1. If ϕ > 1, the sign sequence is *S*_*q*_ = (− ± − +) and the number of sign variations is *V*_*q*_ = 1 or 3. Thus, equation *q*(*y*) = 0 admits 1 or 3 positive roots. Since 1 or 3 of such roots are non-feasible, we have 0 or 2 feasible roots. Hence, only scenarios ø and II are possible if ϕ > 1. We numerically verified the occurrence of scenarios I, and ø and II, under the respective conditions above (Table 2).

**Analysis for** *f* = *f*_*B*_ **and** *g* = *g*_2_. In this case, *q* is a third degree polynomial with coefficients

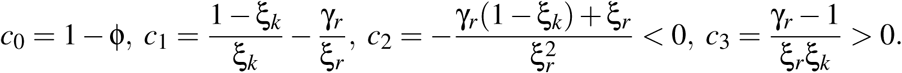

By *lemma 2*, the coefficients *d*_0_ and *d*_3_ of *r*(*u*) are

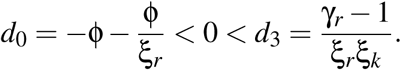

Thus, it follows that 1 or 3 positive roots of *q* satisfy *y* > 1. Now we analyze the sign sequence of *q*. If ϕ < 1, we have *S*_*q*_ = (+ ± − +) and *V*_*q*_ = 2. Thus, *q* admits 0 or 2 positive roots. As at least one satisfies *y* > 1, we conclude that *q* has 2 positive roots but an unique root in the interval 0 < *y* < 1. Hence, only scenario I is possible if ϕ < 1. If ϕ > 1, we have *S*_*q*_ = (− ± − +) and *V*_*q*_ = 1 or 3. Thus, *q* admits 1 or 3 positive roots. Since 1 or 3 of such roots are non-feasible, we have 0 or 2 feasible roots. Hence, only scenarios ø and II are possible if ϕ > 1. We numerically verified the occurrence of scenarios I, and ø and II, under the respective conditions above (see Table 2).

**Analysis for** *f* = *f*_*B*_ **and** *g* = *g*_3_. In this case, *q* is a fourth degree polynomial with coefficients

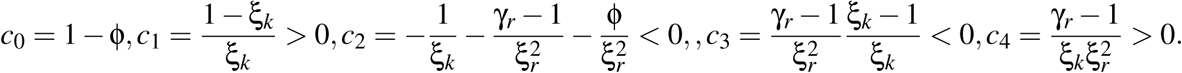

The coefficients *d*_0_ and *d*_4_ of *r*(*u*) are

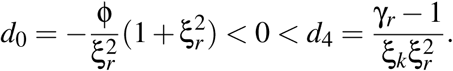

Thus, by *lemma 2*, 1 or 3 positive roots of *q* satisfy *y* > 1. Let us analyse the sign sequence of *q*. If ϕ < 1, then *S*_*q*_ = (++ − − +) and *V*_*q*_ = 2. Thus, *q* admits 0 or 2 positive roots. As at least one satisfies *y* > 1, it follows that *q* has 2 positive roots but an unique root in the interval 0 < *y* < 1. Hence, only scenario I is possible if ϕ < 1. If ϕ > 1, then *S*_*q*_ = (− + − − +) and *V*_*q*_ = 3. Thus, *q* admits 1 or 3 positive roots. Since 1 or 3 of such roots satisfy *y* > 1, we have 0 or 2 feasible roots. Hence, only scenarios ø and II are possible if ϕ > 1. We numerically verified the occurrence of scenarios I, and ø and II, under the respective conditions above (see Table 2).

**Analysis for** *f* = *f*_*B*_ **and** *g* = *g*_4_. In this case, *q* is a fourth degree polynomial. A priori, *q* may admits up to 4 positive roots and scenarios ø, I, II, III and IV would be possible, while scenario V is not possible, since it corresponds to five positive roots for *q*. Indeed, we numerically verified the occurrence of all these scenarios (see Table 2).

**Analysis for** *f* = *f*_*B*_ **and** *g* = *g*_5_. In this case, the next *lemma* guarantees that equation *h*_1_ = *h*_2_ does not admit more than two different positive roots. This implies that only scenarios ø, I and II are possible. We numerically verified the occurrence of these scenarios (see Table 2).

**Lemma 3.** *If* (*f*, *g*) = (*f*_*B*_, *g*_5_), *then equation h*_1_ = *h*_2_ *admits at most two different positive roots.*

*Proof.* Suppose by contradiction that *h*_1_(*y*) = *h*_2_(*y*) admits three or more different positive roots. Since *h*_1_ and *h*_2_ are differentiable functions for all *y* ≥ 0, it follows from the *Rolle theorem* that the difference of their derivatives, 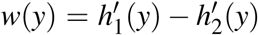, has at least two different positive roots, say 0 < *ŷ*_1_ < *ŷ*_2_. The expressions for 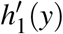 and 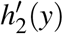 are

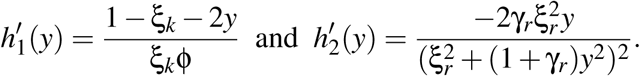

Thus, lim_*y*→∞_ *w*(*y*) = − ∞, which implies that there exists *M* > 0 such that *w*(*y*) < 0 for all *y* ≥ *M*. Hence, *ŷ*_1_ and *ŷ*_2_ are inside the interval (0, *M*). From the *Rolle theorem*, it follows that *w*′ (*y*) has a root 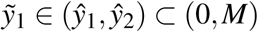.

We shall show that *w*′ (*y*) admits a second root 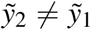 within (0, *M*). If *w*′ (*ŷ*_*i*_) = 0 for some *i*_0_ ∈ {1, 2 }, then just take 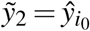. Now, suppose *w*′ (*ŷ*_*i*_) ≠ 0 for *i* = 1, 2. Since *w*(0) = (1 − ξ_*k*_)*/*(ξ_*k*_ϕ) > 0 > *w*(*M*), we conclude that *w*(*y*) changes its sign an odd number of times within the interval [0, *M*]. As *w*′ (*ŷ*_*i*_) = 0 for *i* = 1, 2, it follows that *w*(*y*) changes it sign at the roots *ŷ*_*i*_ and necessarily *w*(*y*) admits a third root *ŷ*_3_ ∈ [0, *M*], distinct from *ŷ*_1_ and *ŷ*_2_. Suppose without loss of generality that *ŷ*_3_ > *ŷ*_2_. Thus, it follows from the *Rolle theorem* that *w*′ (*y*) has another root 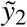 within the interval (0, *M*). Therefore, we’ve proved that 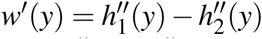 has two distinct roots 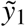 and 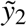 within the interval (0, *M*).

Thus, the curves of 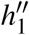 and 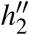 have at least two intersections for *y* > 0. Since

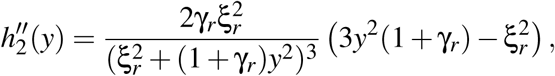

the interval of positive values of *y* where 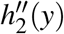 is negative is *I*_*_ = [0, *u*_1_), where 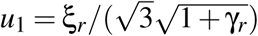. On the other hand, we have that 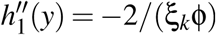 is negative for *y* ≥ 0. Thus, the two roots of 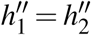 should occur within *I*_*_. The derivative of 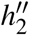,

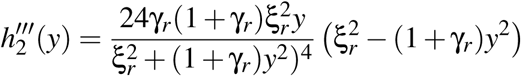

has the following properties: if 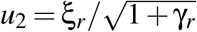, then 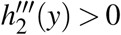 for 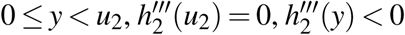 for *y* > *u*_2_, and *u*_2_ > *u*_1_. Thus, 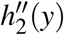 is an increasing function within the interval *I*_*_, while 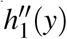 is constant. Therefore, the curves of 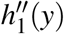 and 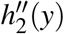 do not have two intersections for *y* > 0, which is a contradiction. Therefore, the equation *h*_1_(*y*) = *h*_2_(*y*) do not have more than two positive roots.

**Analysis for** *f* = *f*_*C*_ **and** *g* = *g*_1_. In this case, *q* is a fourth degree polynomial with coefficients

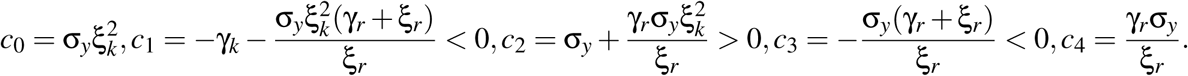

Thus, *S*_*q*_ = (+ − + − +), *V*_*q*_ = 4 and *q* admits 0 or 2 or 4 positive roots. The coefficients *d*_0_ and *d*_4_ or *r* are

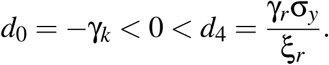

By *lemma 2*, 1 or 3 positive roots of *q* satisfy *y* > 1. Therefore, *q* admits 1 or 3 feasible roots. Hence, only scenarios I and III are possible. We numerically verified the occurrence of such scenarios (see Table 3).

**Analysis for** *f* = *f*_*C*_ **and** *g* = *g*_2_. In this case, *q* is a fourth degree polynomial with coefficients

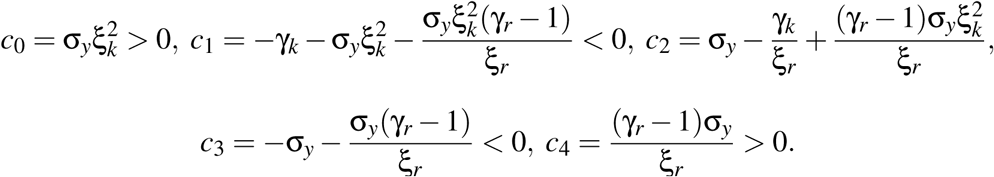

Thus, *S*_*q*_ = (+ − ± − +), *V*_*q*_ = 2 or 4, and *q* admits 0 or 2 or 4 positive roots. The coefficients *d*_0_ and *d*_4_ are

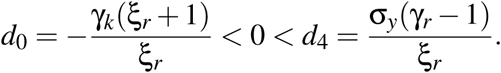

By *lemma 2*, 1 or 3 positive roots of *q* satisfy *y* > 1. Therefore, *q* admits 1 or 3 feasible roots. Hence, only scenarios I and III are possible. We numerically verified the occurrence of such scenarios (see Table 3).

**Analysis for** *f* = *f*_*C*_ **and** *g* = *g*_3_. In this case, *q* is a fifth degree polynomial with coefficients

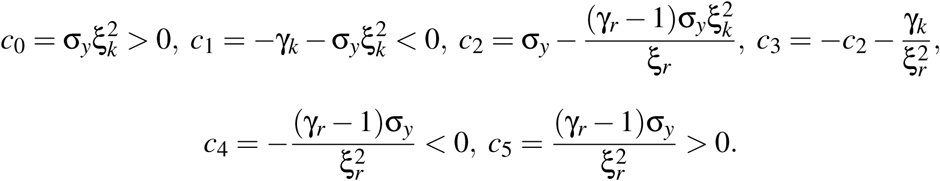

If *c*_2_ > 0, then *c*_3_ < 0, *S*_*q*_ = (+ − + − − +), *V*_*q*_ = 4, and *q* admits 0 or 2 or 4 positive roots. If *c*_2_ < 0, then *S*_*q*_ = (+ − − ± − +), *V*_*q*_ = 2 or 4, and *q* admits 0 or 2 or 4 positive roots. Thus, both cases result in the same possibilities. The coefficients *d*_0_ and *d*_5_ of *r*(*u*) are

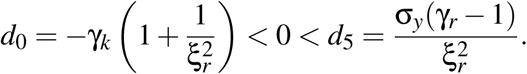

By *lemma 2, q* has 1 or 3 or 5 roots *y* > 1. Therefore, *q* admits 1 or 3 feasible roots. Hence, only scenarios I and III are possible. We numerically verified the occurrence of such scenarios (see Table 3).

**Analysis for** *f* = *f*_*C*_ **and** *g* = *g*_4_. In this case, *q* is a fifth degree polynomial with coefficients

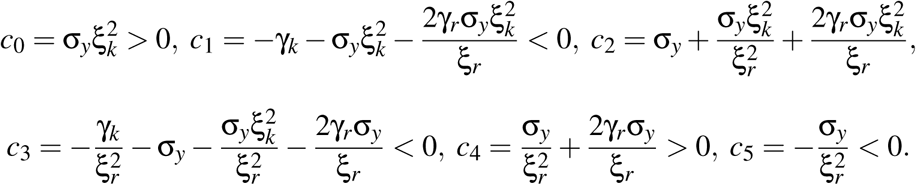

Thus, *S*_*q*_ = (+ − + − + −), *V*_*q*_ = 5, and *q* admits 1 or 3 or 5 positive roots. Hence, only scenarios I, III and V are possible. We numerically verified the occurrence of such scenarios (see Table 3).

**Analysis for** *f* = *f*_*C*_ **and** *g* = *g*_5_. In this case, *q* is a fifth degree polynomial with coefficients

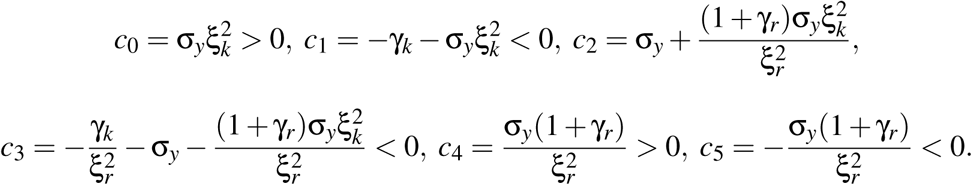

Thus, *S*_*q*_ = (+ − + − + −), *V*_*q*_ = 5, and *q* admits 1 or 3 or 5 positive roots. Hence, only scenarios I, III and V are possible. We numerically verified the occurrence of such scenarios (see Table 3).

**Analysis for** *f* = *f*_*D*_ **and** *g* = *g*_1_. In this case, *q* is a fourth degree polynomial with coefficients

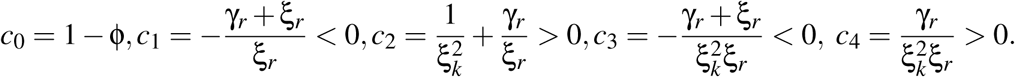

The coefficients *d*_0_ and *d*_4_ of *r*(*u*) are

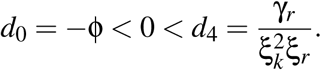

By *lemma 2, q* has 1 or 3 positive roots which satisfy *y* > 1. Analyzing the sign sequence of *q*, we conclude the following. If ϕ < 1, then *S*_*q*_ = (+ − + − +), *V*_*q*_ = 4, and *q* admits 0 or 2 or 4 positive roots. Since 1 or 3 are non-feasible, it follows that *q* has 1 or 3 feasible roots and only scenarios I and III are possible in this case. If ϕ > 1, then *S*_*q*_ = (− − + − +), *V*_*q*_ = 3, and *q* admits 1 or 3 positive roots. Since 1 or 3 are non-feasible, it follows that *q* has 0 or 2 feasible roots and only scenarios ø and II are possible in this case. We numerically verified the occurrence of scenarios ø, I, II and III (see Table 3).

**Analysis for** *f* = *f*_*D*_ **and** *g* = *g*_2_. In this case, *q* is a fourth degree polynomial with coefficients

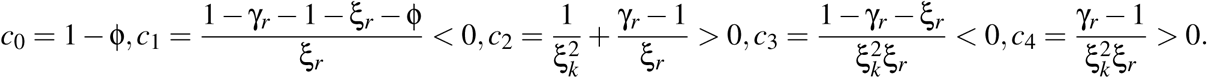

The coefficients *d*_0_ and *d*_4_ of *r*(*u*) are

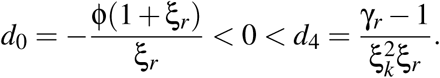

By *lemma 2, q* has 1 or 3 positive roots which satisfy *y* > 1. Analyzing the sign sequence of *q*, we conclude the following. If ϕ < 1, then *S*_*q*_ = (+ − + − +), *V*_*q*_ = 4, and *q* admits 0 or 2 or 4 positive roots. Since 1 or 3 are non-feasible, it follows that *q* has 1 or 3 feasible roots and only scenarios I and III are possible in this case. If ϕ > 1, then *S*_*q*_ = (− − + − +), *V*_*q*_ = 3, and *q* admits 1 or 3 positive roots. Since 1 or 3 are non-feasible, it follows that *q* has 0 or 2 feasible roots and only scenarios ø and II are possible in this case. We numerically verified the occurrence of scenarios ø, I, II and III (see Table 3).

**Analysis for** *f* = *f*_*D*_ **and** *g* = *g*_3_. In this case, *q* is a fifth degree polynomial with coefficients

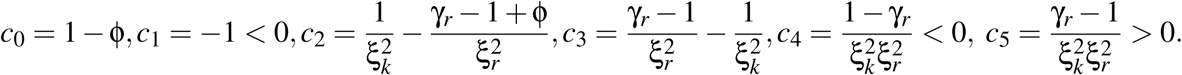

The coefficients *d*_0_ and *d*_5_ of *r*(*u*) are

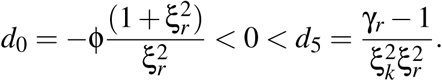

By *lemma 2, q* has 1 or 3 positive roots which satisfy *y* > 1. Analyzing the sign sequence of *q*, we conclude the following. If ϕ < 1, then *S*_*q*_ = (+ − ± ± − +), *V*_*q*_ = 2 or 4, and *q* admits 0 or 2 or 4 positive roots. Since 1 or 3 are non-feasible, it follows that *q* has 1 or 3 feasible roots and only scenarios I and III are possible in this case. If ϕ > 1, then *S*_*q*_ = (− − ± ± − +), *V*_*q*_ = 1 or 3, and *q* admits 1 or 3 positive roots. Since 1 or 3 are non-feasible, it follows that *q* has 0 or 2 feasible roots and only scenarios ø and II are possible in this case. We numerically verified the occurrence of scenarios ø, I, II, and III (see Table 3).

**Analysis for** *f* = *f*_*D*_ **and** *g* = *g*_4_. In this case, *q* is a fifth degree polynomial and all scenarios are possible. We numerically verified the occurrence of each scenario (see Table 3).

**Analysis for** *f* = *f*_*D*_ **and** *g* = *g*_5_. In this case, *q* is a fifth degree polynomial and all scenarios are possible. We numerically verified the occurrence of scenarios ø, I, II, III and IV (see Table 3), while scenario V is excluded due to the following *lemma*.

**Lemma 4.** *If* (*f*, *g*) = (*f*_*D*_, *g*_5_), *then q*(*y*) *does not admit five different roots in the interval y* ∈ [0, 1].

*Proof.* For *f* = *f*_*D*_ and *g* = *g*_5_ we have

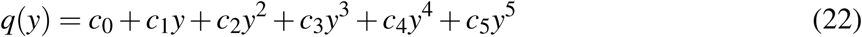

with coefficients

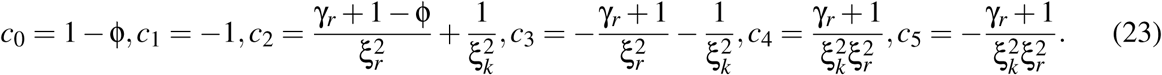

If ϕ > 1, then the sign sequence of *q* is (− −+ − + -), and thus *V*_*q*_ = 4. Hence, *q*(*y*) has at most 4 positive roots and our claim is valid.

Now, assume that ϕ < 1, and suppose by contradiction that *q*(*y*) has 5 different roots within the interval [0, 1]. Then, by the *Rolle theorem*, it follows that *q*′ (*y*) has 4 roots within (0, 1) and then *q*″ (*y*) has three roots within (0, 1). The function *q*″ (*y*) is a third degree polynomial, and we will count its roots by calculating its Sturm sequence and applying the Sturm’s theorem (see Basu et al. [2007], section 2.2.2).

The Sturm sequence of a polynomial *S*(*y*) is a sequence (*S*_0_, *S*_1_, *S*_2_, …) of polynomials defined as follows:

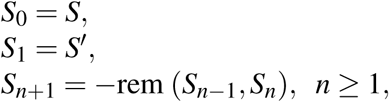

where rem(*S*_*n-*1_, *S*_*n*_) is the remainder of the Euclidean division of *S*_*n-*1_ by *S*_*n*_ (see Basu et al. [2007], section 2.2.2). The number of sign variations of the Sturm sequence evaluated at *c* ∈ℝ, is denoted by *V*_*S*_(*c*) and defined as the number of sign variations in the sequence of numbers

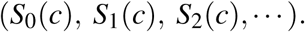

Let *a, b* ∈ℝ. The Sturm‘s Theorem states that the number of roots of *S*(*y*) within the interval [*a, b*] is equal to the difference *V*_*S*_(*a*) − *V*_*S*_(*b*).

We will apply the Sturm‘s Theorem to *S*(*y*) = *q*″ (*y*) and count its roots inside the interval [*a, b*] = [0, 1]. The Sturm sequence of a third degree polynomial

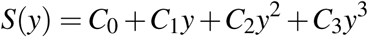

is given by

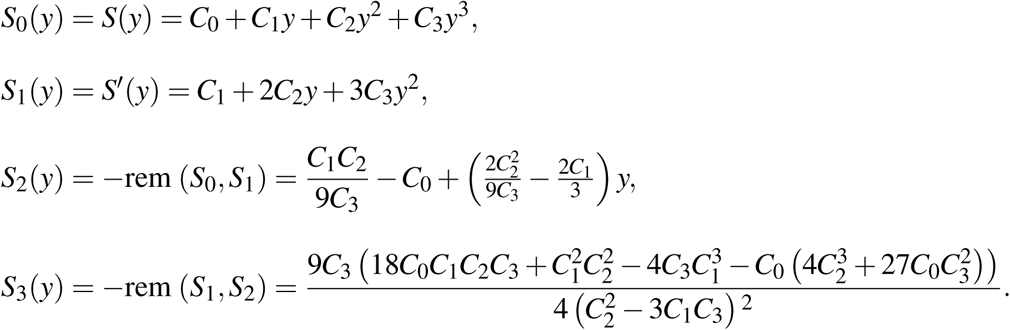

Thus, the Sturm sequence of *S* evaluated at *y* = 0 is

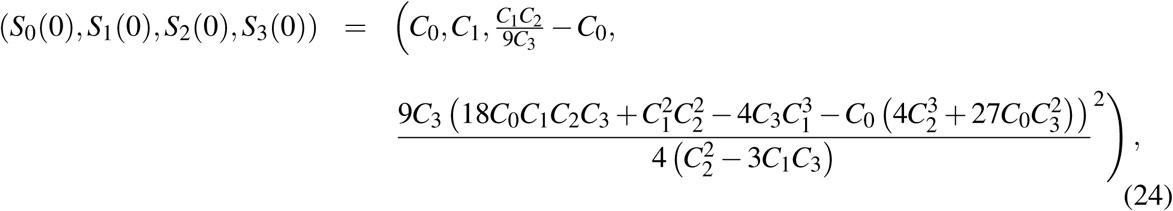

and the Sturm sequence of *S* evaluated at *y* = 1 is

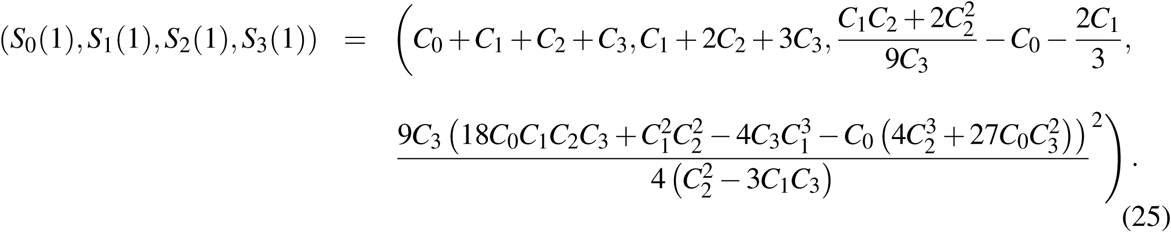

From (22), we have *q*″ (*y*) = 2*c*_2_ + 6*c*_3_*y* + 12*c*_4_*y*^2^ + 20*c*_5_*y*^3^ with *c*_*i*_ given in (23). Writing *S*(*y*) = *q*″ (*y*) = *C*_0_ + *C*_1_*y* + *C*_2_*y*^2^ + *C*_3_*y*^3^, we obtain the coefficients

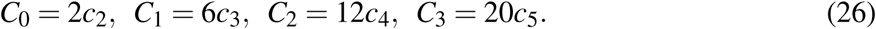

Hence, using formulas (24), (26) and (23), and the fact that ϕ < 1 < γ_*r*_, we obtain that the first three terms of the Sturm sequence of *q*″ evaluated at *y* = 0 are

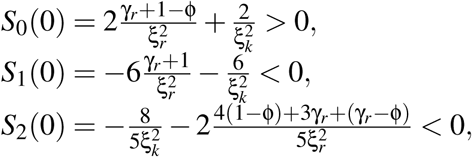

while *S*_3_(0) has a complicated expression with an undetermined sign. However, we can conclude that 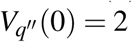 and 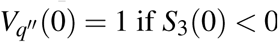.

Now, again using the fact that ϕ < 1 < γ_*r*_ and formulas (25), (26) and (23), we obtain that the first three terms of the Sturm sequence of *q* ″ evaluated at *y* = 1 are

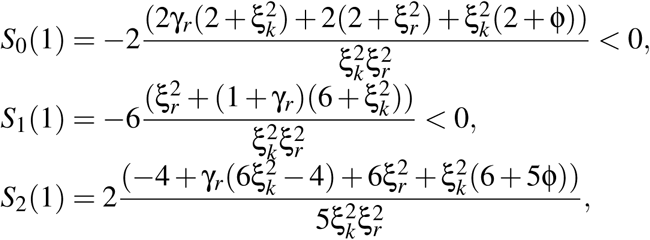

which has an undetermined sign, and *S*_3_(1), which has a complicated expression with an undetermined sign. However, notice from (24) and (25) that *S*_3_(1) = *S*_3_(0). Thus, we can conclude that 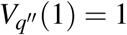 if *S*_3_(0) > 0 and 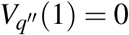 or 2 if *S*_3_(0) < 0.

Therefore, the number of roots of *q*″ (*y*) within the interval [0, 1], which, by the Sturm‘s Theorem, is equal to the difference 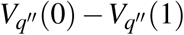, is always less than 3, which is a contradiction. Hence, the equation *q*(*y*) cannot have five roots within the interval [0, 1].

## B Model parameterization and parameter estimation

Here we report the parameter values used for the model simulations. To describe the different scenarios with each sub-model (Tables 2 and 3) we started with the following set of basic values. Parameters regarding proliferating and quiescent LSCs were set to values used in a previous publication and correspond to the median values of a cohort of 122 CML patients [Fassoni et al., 2018]. The values are *p*_*XY*_ = 0.05, *p*_*YX*_ = 0.001, *p*_*Y*_ = 0.2 and *T*_*Y*_ = 10^6^. For immune cells we adopted the values *d*_*Z*_ = 1 month^*−*1^ and *p*_*Z*_ = 10^3^ cells/month so that the normal level of immune cells is *p*_*Z*_*/d*_*Z*_ = 10^3^ cells. Parameters *C*_*K*_ and *C*_*R*_ were set to values defined according to the specific functional responses used in each sub-model. For linear functional responses (*F* = *F*_*A*_ or *G* = *G*_1_), we adopted *C*_*K*_,*C*_*R*_ = 5 *×* 10^5^ meaning that *F* and *G* reach their maximum values for BCR-ABL1/ABL1 ratios around 50%. For the Holling type II and III functional responses (*F* = *F*_*B*_, *F*_*C*_ or *G* = *G*_2_, *G*_3_), we adopted *C*_*K*_,*C*_*R*_ = 10^4^ meaning that *F* and *G* reach half of their maximum values for BCR-ABL1/ABL1 ratios around 1%. For immune window (*F* = *F*_*D*_ or *G* = *G*_4_) and the immune suppression (*G* = *G*_5_) functional responses, we adopted *C*_*K*_,*C*_*R*_ = 10^3^ meaning that *F* and *G* reach their maximum values for BCR-ABL1/ABL1 ratios around 0.1%, i.e. MR3. All these basic values above were used as starting values for searching the possible scenarios for each submodel. Parameters *m*_*K*_ and *m*_*R*_ did not have a specific starting value. By varying the least possible number of parameters (starting with *m*_*K*_, *m*_*R*_, then *C*_*K*_, *C*_*R*_ and then *p*_*Y*_), we obtained the parameter values shown in Tables 2 and 3, leading to different scenarios for each possible sub-model.

To estimate the model parameter values corresponding to the fits in Figure 3, we allowed parameters *p*_*Y*_, *p*_*YX*_, *m*_*K*_ and *m*_*R*_ to vary and used a minimization algorithm to find those parameter values that minimize the quadratic error between the model solution and the patient data, defined as

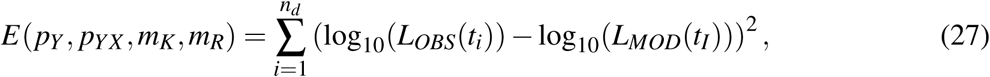

where *L*_*OBS*_(*t*_*i*_) are the observed BCR-ABL1/ABL1 ratios at times *t*_*i*_ in the patient time course with *n*_*d*_ = 30 data-points, and *L*_*MOD*_(*t*_*i*_) = 100*Y* (*t*_*i*_)*/T*_*Y*_ are the simulated BCR-ABL1/ABL1 ratios at the same time points. The following intervals for parameter searches were used: 0.05 ≤ *p*_*Y*_ ≤ 1, 0 ≤ *m*_*K*_ ≤ 1000, *d*_*Z*_ < *m*_*R*_ ≤ 1000, and 0 ≤ *p*_*YX*_ ≤ 0.1. The other parameters remained constant and we assumed the following values: regarding leukemic cells, we used the values obtained in our previous model for a specific patient: *p*_*XY*_ = 0.0451256, *T*_*Y*_ = 10^6^ and *e*_*TKI*_ = 0.493541 + *p*_*Y*_ [Fassoni et al., 2018]. For parameters *d*_*Z*_, *p*_*Z*_, *C*_*K*_, *C*_*R*_ we adopted the same values as above (values for *C*_*K*_ and *C*_*R*_ varied according to each sub-model). The results with the estimated values are given in Table 4.

**Table 4:** Estimated values for *p*_*Y*_, *p*_*YX*_, *m*_*K*_ and *m*_*R*_, used in Figure 3.

| $(F, G)$ | $p_Y$ | $m_K$ | $m_R$ | $p_{YX}$ |
| --- | --- | --- | --- | --- |
| $(F_A, G_1)$ | 0.18939321 | 0.00006908 | 3.31196830 | 0.00298540 |
| $(F_A, G_2)$ | 0.18940069 | 0.00003577 | 1.19091170 | 0.00298540 |
| $(F_A, G_3)$ | 0.18836653 | 0.00002774 | 1.23852800 | 0.00298517 |
| $(F_A, G_4)$ | 0.21386851 | 0.43721542 | 1.22494960 | 0.00315841 |
| $(F_A, G_5)$ | 0.18929618 | 0.00093258 | 9.66089240 | 0.00298538 |
| $(F_B, G_1)$ | 0.18940055 | 0.00002076 | 5.76081590 | 0.00298541 |
| $(F_B, G_2)$ | 0.18940214 | 0.00000225 | 1.29802820 | 0.00298540 |
| $(F_B, G_3)$ | 0.18939853 | 0.00000192 | 1.23391420 | 0.00298540 |
| $(F_B, G_4)$ | 0.21721919 | 0.00597071 | 1.21093450 | 0.00316262 |
| $(F_B, G_5)$ | 0.18929618 | 0.00093258 | 9.66089240 | 0.00298538 |
| $(F_C, G_1)$ | 0.18939538 | 0.00000256 | 6.04955220 | 0.00298540 |
| $(F_C, G_2)$ | 0.18940266 | 0.00000309 | 1.27006090 | 0.00298540 |
| $(F_C, G_3)$ | 0.18939157 | 0.00000223 | 1.25669390 | 0.00298540 |
| $(F_C, G_4)$ | 0.21534823 | 0.01383360 | 1.32215430 | 0.00315400 |
| $(F_C, G_5)$ | 0.18938712 | 0.00000897 | 2.59943410 | 0.00298539 |
| $(F_D, G_1)$ | 0.31487806 | 0.09655284 | 4.78823520 | 0.00390086 |
| $(F_D, G_2)$ | 0.32421736 | 0.11156311 | 1.00156410 | 0.00409459 |
| $(F_D, G_3)$ | 0.30345149 | 0.09890912 | 1.00002180 | 0.00393134 |
| $(F_D, G_4)$ | 0.24723328 | 0.00845071 | 1.00001490 | 0.00338385 |
| $(F_D, G_5)$ | 0.18506692 | 0.00023119 | 94.0396200 | 0.00298504 |

To generate Figure 8, we fixed all parameters, with exception of *m*_*K*_ and *m*_*R*_, which were allowed to vary in order to lead to different outcomes after cessation for each selected sub-model. The values for *m*_*K*_ and *m*_*R*_ were manually selected and are given in Table 5. Regarding the other parameters, in all simulations we used the fixed values *p*_*YX*_ = 0.00353003, *p*_*Y*_ = 0.2, *e*_*TKI*_ = 0.693541, *p*_*XY*_ = 0.0451256, while *T*_*Y*_ *d*_*Z*_, *p*_*Z*_, *C*_*K*_, and *C*_*R*_ are the same as above (values for *C*_*K*_ and *C*_*R*_ varied according to each sub-model).

**Table 5:**
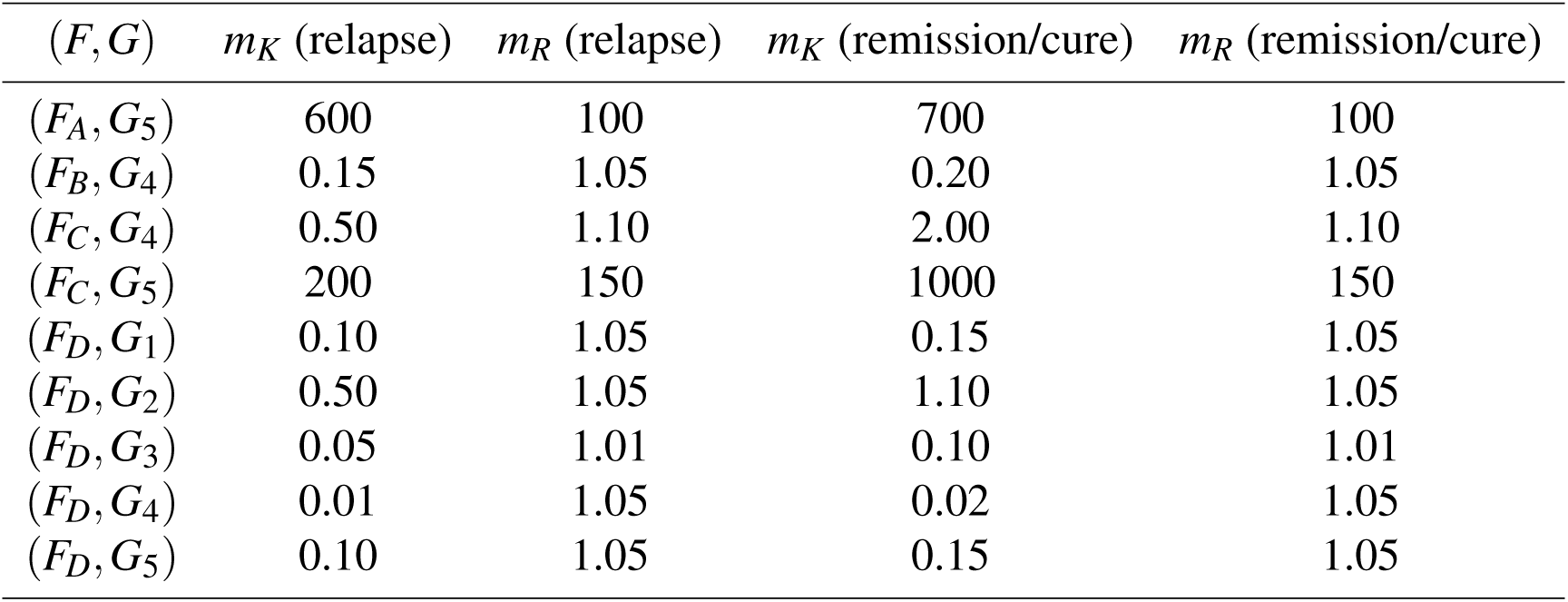
Values for *m*_*K*_ and *m*_*R*_ used to generate the different outcomes in the simulations shown in Figure 8: relapse (solid lines) and remission/cure (dashed lines).

## Acknowledgements

The authors wish to thank Christoph Baldow and Tom Haehnel for their comments and suggestions that contributed to this work.

